# The Th1/Th17 axis regulates chimeric antigen receptor (CAR) T cell therapy toxicities

**DOI:** 10.1101/2025.03.06.641668

**Authors:** Payal Goala, Yongliang Zhang, Nolan Beatty, Shannon McSain, Cooper Sailer, Muhammad Junaid Tariq, Showkat Hamid, Eduardo Cortes Gomez, Jianmin Wang, Justin C Boucher, Constanza Savid Frontera, Sae Bom Lee, Hiroshi Kotani, Michael D. Jain, Marco L. Davila

**Author notes:** Correspondence to Marco L. Davila.

## Abstract

CAR-T therapy has led to significant improvements in patient survival. However, a subset of patients experience high-grade toxicities, including cytokine release syndrome (CRS) and immune cell-associated hematologic toxicity (ICAHT). We utilized IL-2Rα knockout mice to model cytokine toxicities with elevated levels of IL6, IFNγ, and TNFα and increased M1-like macrophages. Onset of CRS was accompanied by a reduction in peripheral blood neutrophils due to disruption of bone marrow neutrophil homeostasis characterized by an increase in apoptotic neutrophils and a decrease in proliferative and mature neutrophils. Both non-tumor-bearing and Eμ-ALL tumor-bearing mice recapitulated the co-occurrence of CRS and neutropenia. IFNγ-blockade alleviated CRS and neutropenia without affecting CAR-T efficacy. Mechanistically, a Th1-Th17 imbalance was observed to drive co-occurrence of CRS and neutropenia in an IFNγ-dependent manner leading to decreased IL-17A and G-CSF, neutrophil production, and neutrophil survival. In patients, we observed an increase in the IFNγ-to-IL-17A ratio in the peripheral blood during high-grade CRS and neutropenia. We have uncovered a biological basis for ICAHT and provide support for the use of IFNγ-blockade to reduce CRS and neutropenia.

**Statement of Significance:** Despite clinical success of CAR-T therapy, patients develop toxicities such as cytokine release syndrome and neutropenia, whose co-occurrence impacts their survival and quality-of-life. We recapitulate these toxicities in mice to discover their co-occurrence is driven by Th1-Th17 imbalance following CAR-T administration, which can be prevented via IFNγ blockade.

## Introduction

CAR-T therapy has improved overall and complete response rates, and prolonged progression free survival (PFS) of patients with relapsed/refractory hematological malignancies [1]. The short-term and long-term impacts of anti-CD19 CAR-T products on the immune landscape of B cell malignancies has garnered considerable attention in recent years [2, 3], most notably pertaining to CAR-T associated toxicities, which include CRS (cytokine release syndrome) and ICANS (Immune cell associated neurotoxicity syndrome). CRS arises from the excessive release of cytokines such as IFNγ and TNFα [4] from CAR-T cells that activate macrophages and other immune cells to release more pro-inflammatory cytokines, which trigger a cascade of events leading to symptoms such as fever, hypotension, hypoxia, and organ toxicities [2]. Overactivation of ANG-2 and Von Willebrand factor during this process causes endothelial dysfunction that is associated with ICANS [5]. Recent clinical studies have shown that CAR-T treated patients also suffer from cytopenias [6]. The mechanisms leading to CAR-T associated cytopenia remains elusive. Rejeski et. al. showed that severe cytopenia in patients post CAR-T treatment led to frequent infections and increased hospitalizations [7, 8]. Onset of cytopenia was associated with worse survival of patients post CAR-T treatment thereby emphasizing the need to address these hematological toxicities [6]. An observation drawn from two independent clinical trials was the co-occurrence of cytopenia with CRS [6, 9]. Among the cytopenias neutropenia was most predominant [6], where the decline in neutrophils intensified with the grade of CRS [7, 9]. These prolonged cytopenias have been labelled as immune cell-associated hematologic toxicity (ICAHT) and are a substantial cause of non-relapse mortality (NRM), but with a lack of effective treatments and little known about their underlying cause [10–12]

High disease burden and excess CAR-T expansion are major factors leading to inflammation that drives CAR-T toxicities [2, 13]. Fischer et. al. showed that multiple myeloma (MM) patients with a smaller proportion (among T cells) of CD25^+^CD127^low^ Tregs (Regulatory T cells) at lymphodepletion were associated with improved response to CAR-T treatment, who also had a higher maximum CRP (C-reactive protein, an indicator of CRS) [14]. Firestone et. al. noted that MM patients who experienced CRS following treatment with a bispecific T cell engager had significantly longer PFS, where a smaller pre-treatment Treg (CD25^+^CD127^low^) proportion among T cells (and an increase in CD8+ T cell proportion) was found to be one of the major drivers for improved response [15]. Kitamura et. al. confirmed a direct correlation between onset of CRS and a significant decrease in Tregs (CD4^+^CD25^+^CD127^-/low^ T cells) [16]. These clinical findings suggest an association between CRS, response to T cell therapies, and Tregs (CD25^+^CD127^low^). IL-2Rα (also known as CD25), a key mediator of lymphoproliferation, is essential for Treg expansion and survival [17, 18] so we used an *in vivo* IL-2Rα knockout (-/-) mice to model toxicities during CAR-T treatment.

## Results

### IL-2Rα-/- Mice Recapitulate CRS Following CAR-T Administration

Two important factors affecting the severity of CRS in patients post CAR-T therapy are extent of CAR-T proliferation and the release of pro-inflammatory cytokines [3]. We hypothesized that knockout of IL-2Rα can emulate the pro-inflammatory environment during CRS. We inoculated IL2Rα-/- (KO) mice as well as WT control mice with the syngeneic B cell leukemia cell line, Eμ-ALL [19]. After a week from tumor engraftment KO and WT mice were administered cyclophosphamide and a day later CD1928ζ CAR-T cells (Fig. 1 A). CAR-T expansion was confirmed in IL-2Rα-/- mice by the presence of GFP+ CAR-T cells in spleen and lymph node. The extent of CAR-T infiltration was significantly higher in KO mice compared to WT and this correlated with greater B cell (B220+) aplasia (Fig. 1 B – C). Enhanced CAR-T expansion was also observed in peripheral blood of KO compared to WT mice, which led to improved efficacy in B cell killing (Fig. 1 D – E). KO mice treated with CAR-T displayed poor survival of less than 35 days (p<0.0001) compared to WT mice that survived more than 100 days (Fig. 1 F). Laboratory analysis demonstrated these deaths were due to CRS and not progressive cancer. We identified infiltration of leukocytes and histiocytes in the H & E-stained sections of splenic red pulp, lymph node, liver, and lung tissues (Fig. 1 G). Evaluation of serum cytokines at week 1 post CAR-T administration in KO mice confirmed the significant elevation of pro-inflammatory cytokines IL-6, TNFα and IFNγ (Fig. 1 H, J, L). There were no notable differences in serum cytokines of CAR-T treated WT mice (Fig. 1 I, K, M).

**Figure 1:**
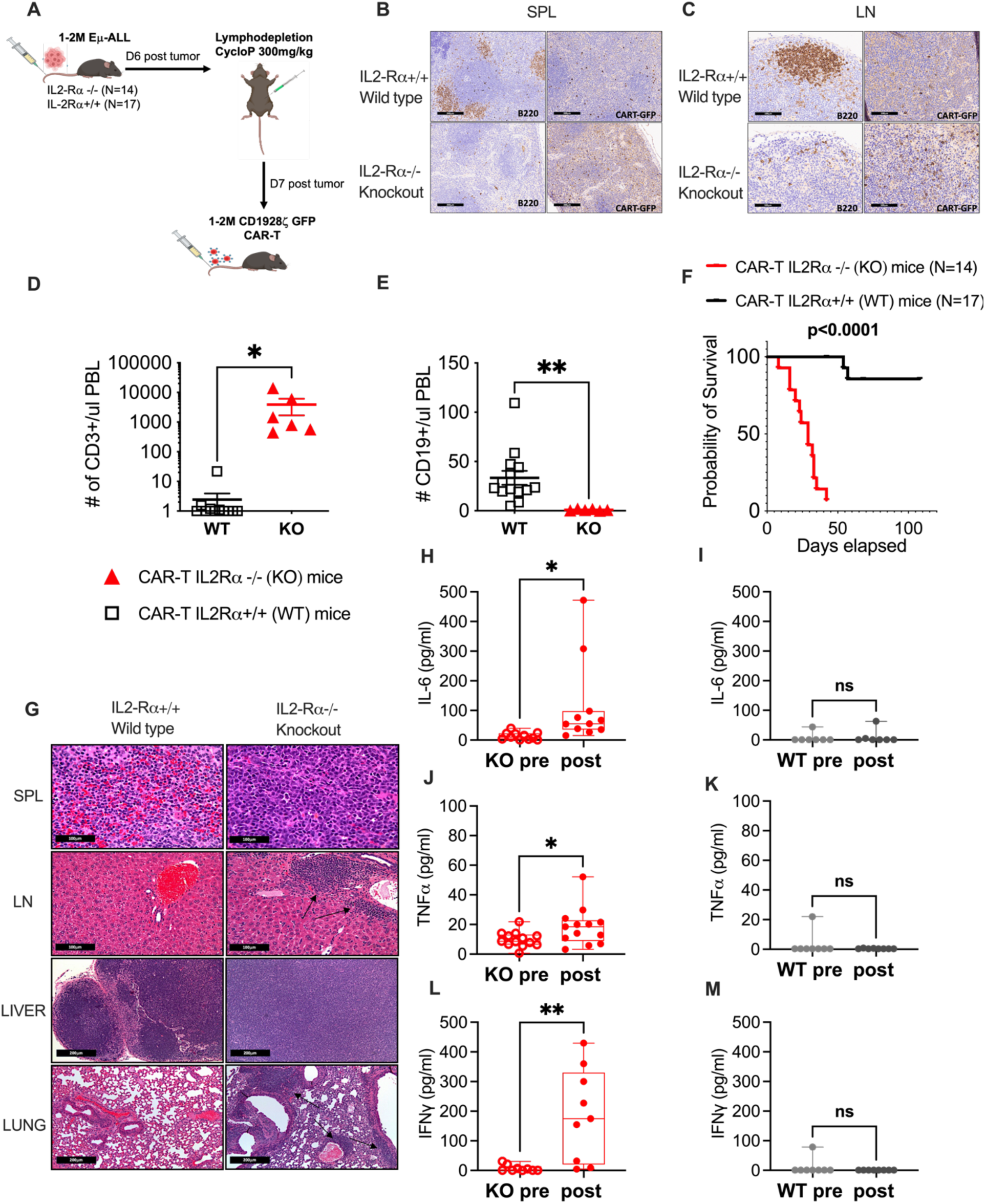
IL2Rα-/- mice recapitulate CRS following CAR-T administration. A. Schematic representation of IL-2Rα-/- (KO) or wild type (WT) mice inoculated with 1 to 2 Million (M) Eμ-ALL cells followed by 300mg/kg cyclophosphamide (CycloP) and 1 to 2M CD1928ζ GFP CAR-T infusion. Data is pooled from 2 independently performed experiments. B. Histopathology sections of Spleen (SPL) at 100x (black scale bar:200μm) and C. Lymph node (LN) at 400X (black scale bar:100μm) a week post CAR-T infiltration in IL-2Rα-/- (KO) vs control (WT) mice. The panel to the left shows B220+ cells (brown) and panel to the right shows CAR-T cells (brown). D. % CD3+ and E. % CD19+ cells in peripheral blood (PBL) were compared between a subset of IL2Rα-/- (KO, N=6) and wild type (WT, N=14) mice at week 1 post CAR-T. F. Kaplan-Meier overall survival curves of KO vs WT mice treated with CD1928ζ CAR-T cells. G. H& E-stained sections of (Top to Bottom) SPL, LN (at 400x; black scale bar:100μm), Liver and lung (at 100x; black scale bar:200μm) from KO and WT mice at week 1 post CAR-T infusion. Arrows shown in LN and lung sections in E represent a mixed leukocyte infiltration including large histocytic cells. Week 1 serum cytokine analysis of IL-6 (H, I), TNFα (J, K) and IFNγ (L, M) in CAR-T treated KO and WT mice respectively. In I, K, M, cytokines were measured for 7 WT mice, while in H, J, L, cytokines were measured for all KO mice that had samples for both pre and post cytokines as paired values from each mouse. Error bars represent standard error of mean (SEM). P values *P < .05, **P < .01, and ***P < .001 were considered significant. P values for cytokine bar plots D-E and H-M were generated using paired t test. P values for Kaplan-Meier survival curve F was generated using Log-rank (Mantel-Cox) test.

### Reversal of CRS Phenotype in IL-2Rα-/- Mice via IL-6-blockade, IFNγ-blockade, and Treg Adoptive Transfer

While lymphodepletion followed by CAR-T treatment best reflects clinical practice, it also confounds the role of CAR-T on the endogenous immune landscape. Therefore, we treated KO and WT (C57BL/6) mice with CAR-T cells only, i.e. without any tumors or lymphodepleting agent cyclophosphamide (Fig. 2 A). A significant elevation of CRS associated pro-inflammatory cytokines IL-6 (p=0.003), TNFα (p=0.041) and IFNγ (p=0.027) were observed after four weeks of CAR-T treatment (Fig. 2 B - D). To validate CAR-T associated CRS, KO mice were treated with either anti-IL-6R monoclonal antibody (mAb) or anti-IFNγ mAb, which are equivalents for Tocilizumab and Emapalumab, respectively. Tocilizumab is used to manage CRS in a clinical setting, whereas Emapalumab has recently been shown to mitigate CRS also [20]. Both anti-IL-6R and anti-IFNγ monoclonal antibodies were administered twice every week, followed by weekly collection of peripheral blood for complete blood profiling and serum analysis until endpoint. Compared to CAR-T only treatment we report baseline restoration of cytokines TNFα, whereas IL-6 and IFNγ levels remained high in KO mice post treatment with anti-IL-6R mAb (Fig. 2 E - G). Treatment with anti-IFNγ mAb (Fig. 2 H - J) reduced IL-6 and TNFα but again noted an increase in IFNγ. Increase in targeted cytokines may be due to drug physiology or secretion by other immune cells. For example, increase in IL-6 (Fig. 2 E) can be due to receptor shedding in response to the action of anti-IL-6R mAb [21, 22]. Similarly, increase in IFNγ (Fig. 2 J) following treatment with anti-IFNγ mAb can be a consequence of target-mediated drug disposition (TMDD) [23–25]. To evaluate this we measured CXCL9, which is induced by IFNγ and an established marker for functional measurement of IFNγ neutralization [23–25].While CAR-T treatment showed a significant increase in CXCL9 (p=0.007), administration of anti-IFNγ mAb led to a significant inhibition of CXCL9 (p<0.0001), thereby confirming the abatement of IFNγ signaling (Fig. 2 K – N). A six-week survival analysis of KO mice (Fig. 2 A) treated with CAR-T showed a significant decrease in survival compared to anti-IL-6R mAb (p=0.022) and anti-IFNγ mAb (p=0.0001) groups (Fig. 2 O).

**Figure 2:**
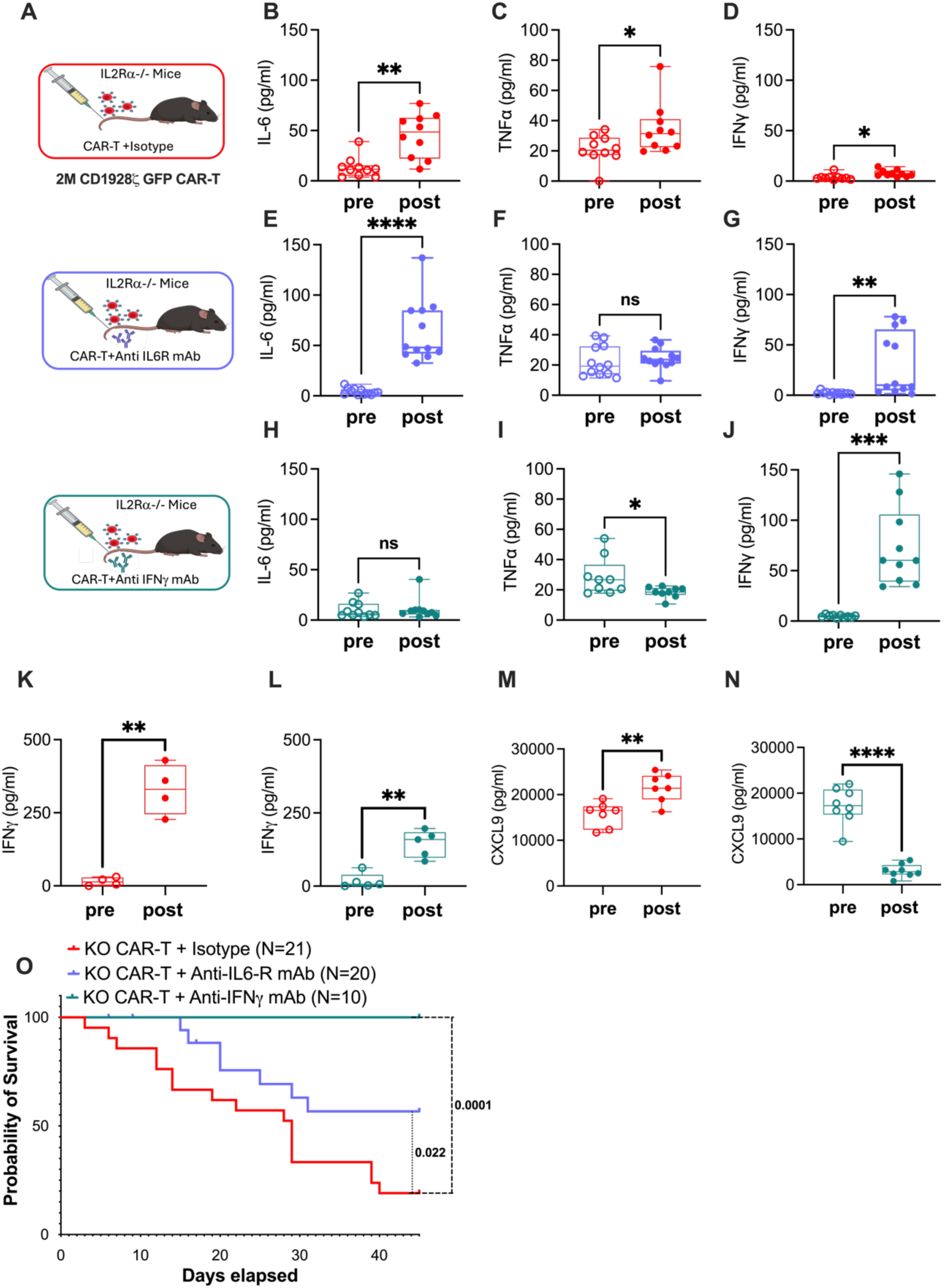
Mitigating CRS in IL2Rα-/- mice. A. Schematic representation of 51 IL2Rα-/- (KO) mice treated with 2M CAR-T followed by administration of either Isotype IgG or 12.5 mg/kg (i.p.) of anti-IL-6R mAb or anti-IFNγ mAb on a twice weekly basis. Data is pooled from 2 independantly performed experiments. B-J. Serum analysis in pre vs week 4 post CAR-T treated mice of cytokines IL-6 (B, E, H), TNFα (C, F, I) and IFNγ (D, G, J). Pre and post cytokines are paired values taken from each mouse alive at week 4 for each cohort (N=10 for CAR-T +Isotype, N=12 for CAR-T + anti-IL-6R mAb and N=10 for CAR-T + anti-IFNγ mAb). K-N. Comparison of IFNγ (K, L) and CXCL9 (M, N) between CAR-T treated KO mice with or without anti-IFNγ mAb. Pre and post cytokine values from K-N are paired values taken from KO mice that had samples for both pre and post cytokines at week 1 (N=7 for CAR-T Group and N=8 for CAR-T + anti-IFNγ mAb group). Data from K-N are from one experiment. O. The impact of CRS management in KO mice was determined by measuring Kaplan-Meier overall survival. Error bars represent SEM. P values *P < .05, **P < .01, and ***P < .001 were considered significant. P values for cytokine bar plots B-J and K-N were generated using paired t test. P values for Kaplan-Meier survival curve O was generated using Log-rank (Mantel-Cox) test.

IL-2Rα-/- mice have dysfunctional Tregs and we hypothesized that CRS in KO mice post CAR-T therapy could in part be due to this deficiency. Clinical studies have similarly shown reduced Treg frequency accounted for higher rate of CAR-T associated ICANS [26] and CRS [14, 27]. KO mice were treated with CAR-T alone (Group 1), or in combination with adoptively transferred low dose Tregs (0.25 M; Group 2), or high dose Tregs (1 M; Group 3) infused prior to CAR-T (Fig. S1 A). Lower Treg dosing was better at improving survival of mice (CAR-T+0.25M Tregs vs CAR-T alone: p=0.056) compared to higher Tregs (CAR-T+1M Tregs vs CAR-T alone: p=0.265) (Fig. S1 B). Compared to CAR-T alone (Fig. S1 C – E) low and high dose Tregs reduced IL-6, TNFα and IFNγ to baseline (Fig. S1 F – K).

### IL-2Rα-/- Mice Show Co-occurrence of CRS and Neutropenia Following CAR-T Infusion

Higher incidence of cytopenias are reported in patients who develop CRS [6, 9]. Longitudinal analysis of KO mice compared to WT post CAR-T treatment (Fig. 3 A) showed elevation of IL-6 (p=0.022), TNFα (p=0.024) and IFNγ (p=0.023) (Fig. 3 B – D), while complete blood profiling showed significant (p=0.007) decline in neutrophils and red blood cells (Fig. 3 E,F). However, a consistent pattern of thrombocytopenia could not be confirmed in KO mice (Fig. 3 G). Neither neutropenia, anemia, nor thrombocytopenia were observed in WT mice. To determine the extent of CAR-T associated neutropenia we compared neutrophil levels of KO mice treated with CAR-T alone, or lymphodepletion alone, or in combination (Fig. S2 A). Both CAR-T alone and lymphodepletion alone led to a drop in neutrophils within one week, however lymphodepletion alone showed a rapid recovery in neutrophils within five weeks and CAR-T alone had prolonged neutropenia (p=0.003) (Fig. S2 B). Lymphodepletion followed by CAR-T similarly showed poor recovery in neutrophils (p=0.047).

**Figure 3:**
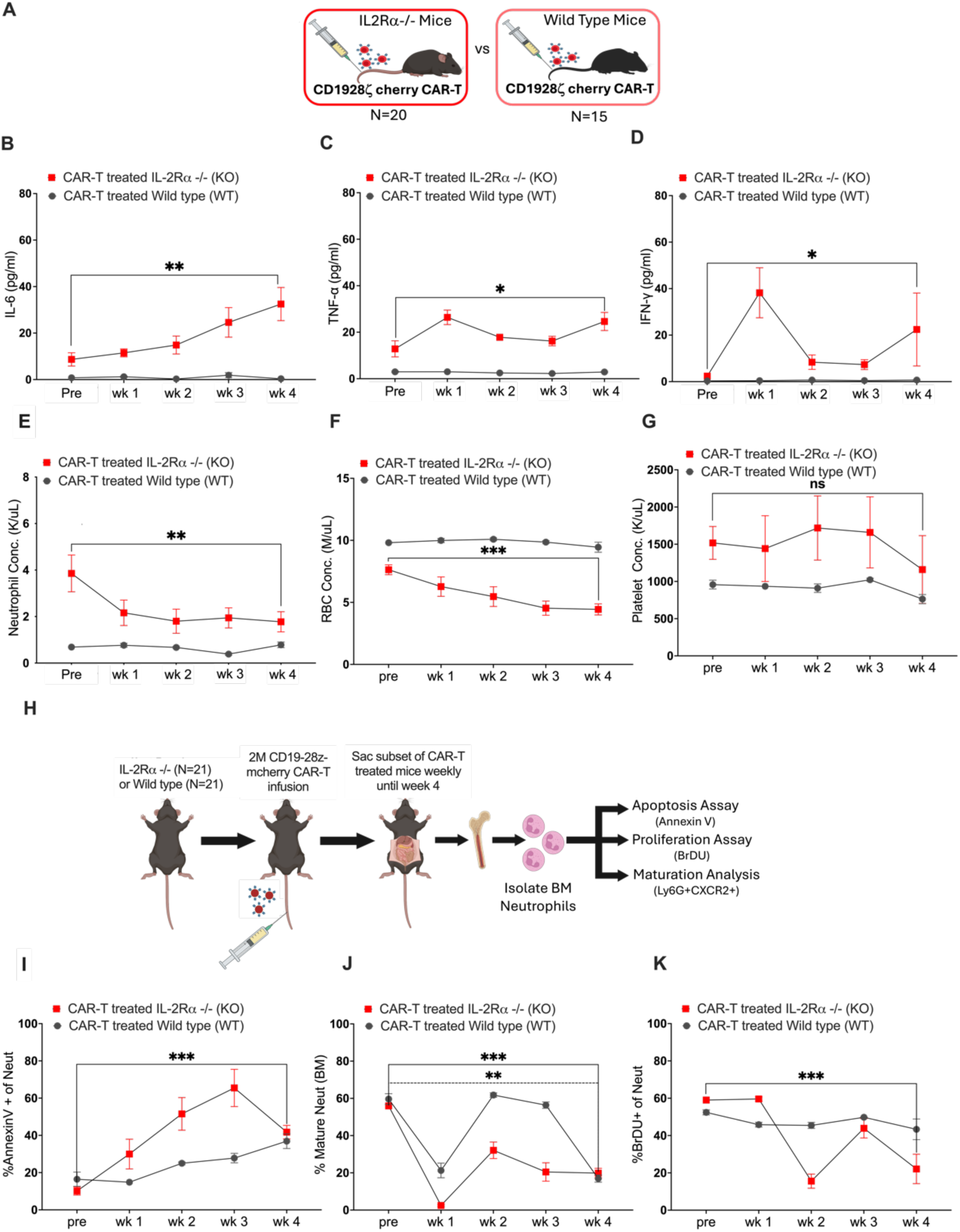
Evaluating co-occurrence of CRS and Neutropenia in IL2Rα-/- mice. A. Schematic representation of IL2Rα-/- (KO) and Wild type (WT) mice treated with 2M CAR-T (i.v.) cells. Data is pooled from 2 independently performed experiments. B-D. Time course analysis of cytokines IL-6 (B), TNFα (C) and IFNγ (D) in post CAR-T treated KO and WT mice. KO pre and post cytokines are paired values taken from each mouse alive until week 4 (N=13). WT paired cytokine values represent a subset of live mice until week 4 (N=7). E-G. Time course of neutrophil concentration (K/μL) (E), red blood cell (RBC) concentration (M/μL) (F) and platelet concentration (K/μL) (G) in the peripheral blood from KO and WT mice using complete blood profiling (CBC). Pre and post levels from CBC are paired values taken from each mouse alive until week 4 per cohort (N=8 for KO and N=8 for WT). H. Schematic of KO and WT mice treated with CAR-T cells sacrificed periodically followed by kit-based isolation of neutrophils from bone marrow (BM). Cell turnover of the purified neutrophils was analysed for apoptosis using Annexin V, and proliferation using BrDU. Neutrophil maturation rate was analysed using BMMC. Data is pooled from 2 independently performed experiments. I-J. Time point comparison in KO vs WT mice of % Annexin V+ apoptotic cells as a frequency of neutrophils (kit-based purification) (I) and % Ly6G+CXCR2+ cells as a frequency of mature neutrophils (gated on BMMC as Live+Lineage-CD11b+Gr1+ckit-CXCR4-cells) (J). (K). KO and WT micewere injected with 2mg BrDU i.p. 48 hours prior to harvest, followed by staining for BrDU labelled neutrophils. Line plot represents % BrDU+ cells as a frequency of live neutrophils (kit-based purification) at weeks 1 through 4. Error bars represent SEM. P values *P < .05, **P < .01, and ***P < .001 were considered significant. P values for line plots B-G were generated using paired t test. P values for line plots I-K were generated using unpaired t test.

Neutrophils are developed in the bone marrow following granulopoiesis [28, 29]. Since neutrophils are short lived a healthy pool of mature (CD11b+Ly6G+CXCR2+) neutrophils in the bone marrow is essential to maintain circulating neutrophils [30]. Additionally, apoptotic neutrophils are homed back to the bone marrow for elimination [31]. To determine the effect of CAR-T cells on neutrophil homeostasis, KO and WT mice injected with CAR-T cells were sacrificed on a weekly basis to harvest bone marrow mononuclear cells (BMMC) (Fig. 3 H). BMMCs collected from KO and WT mice per time point were used for isolating neutrophils stained with Annexin V (marker for apoptosis) as well as BrDU (marker for proliferation). The percentage of total mature neutrophils (characterized as CD11b+Ly6G+CXCR2+ cells) present in the bone marrow pre and post treatment were analyzed by flow cytometry. KO mice showed a significant increase in the rate of apoptosis (p<0.0001) as indicated by an increase in Annexin V+ 7-AAD+ neutrophils, classified as late-stage apoptotic cells that peaked at week 3 post CAR-T (Fig. 3 I). There was no noticeable difference in apoptosis among neutrophils derived from CAR-T treated WT mice. Although a significant decrease in mature neutrophils was observed in both KO and WT (Fig. 3 J) mice, only KO mice had a significant increase in both neutrophil apoptosis and reduced proliferation, confirmed by a decrease (p=0.002) in BrDU+ neutrophils (Fig. 3 K).

### IFNγ-blockade Aids Recovery of CRS and Neutropenia Following CAR-T

Macrophages mediate CRS by releasing pro-inflammatory cytokines [32, 33]. Blockade of IFNγ inhibits macrophage activation caused during CRS ex vivo [20, 34]. Although prevalence of IFNγ has been shown to be less favorable for emergency granulopoiesis [35, 36], the effect of IFNγ blockade on neutropenia in the context of CAR-T associated toxicity has not been characterized. Therefore, KO mice administered with CAR-T cells were divided into three groups: Group 1 included mice treated with CAR-T, Group 2 included mice treated with CAR T cells derived from IFNγ knockout (-/-) mice, and Group 3 mice were treated with CAR-T followed by anti-IFNγ mAb (Fig. 4 A). Week 4 analysis measured CRS associated cytokines (IL-6, TNFα, and IFNγ) and cytokines that regulate neutrophils (IL-17A and G-CSF). There were no observable differences between the activation (CD69 and Granzyme B) and exhaustion (PD1 and TIM3) status of IFNγ-/- CAR-T cells when compared to wild type (Fig. S3 A – E).

**Figure 4:**
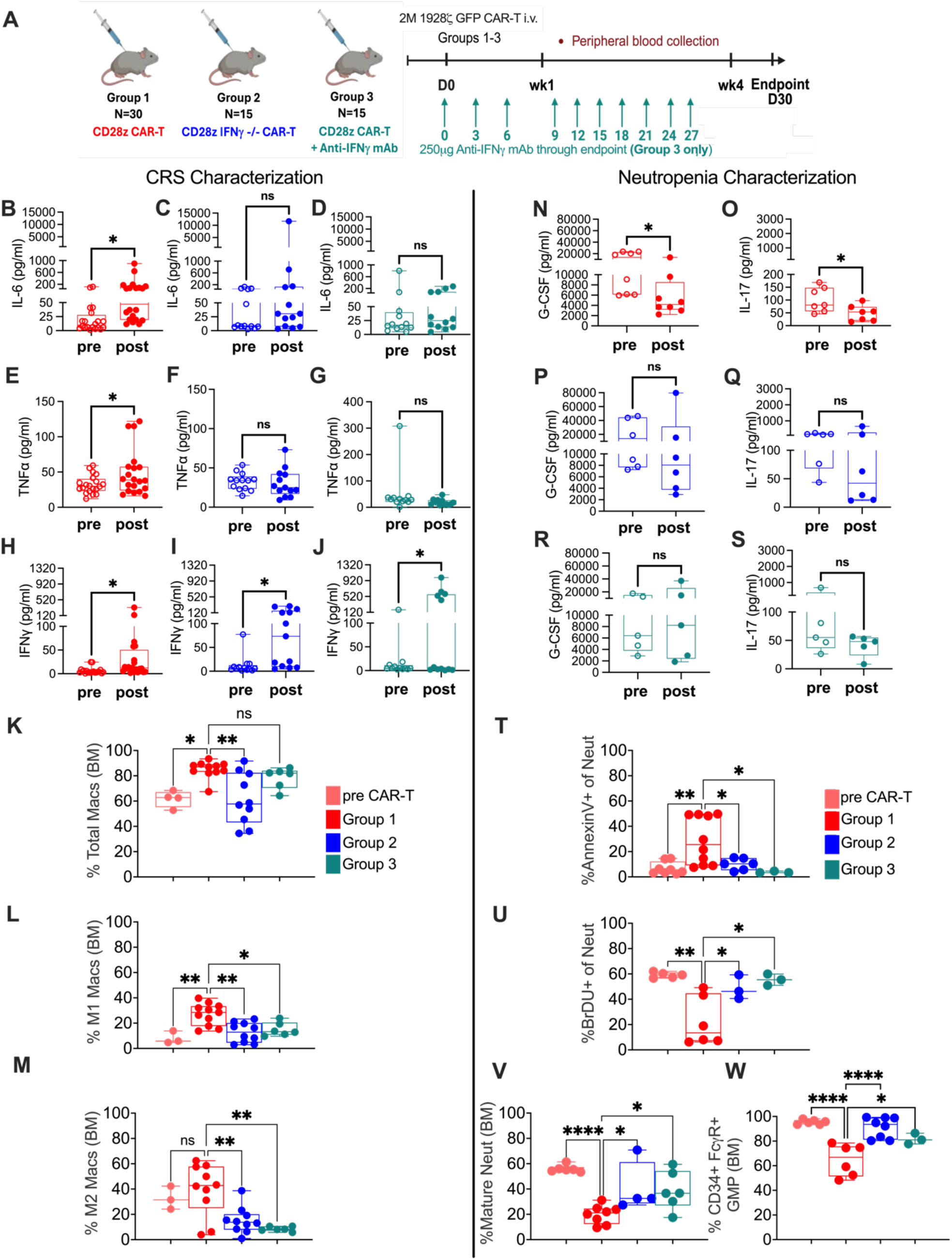
Effect of IFNγ blockade on CRS and Neutrophils. A. KO mice were treated with CAR-T alone (Group 1), or with IFNγ-/- CAR-T cells (Group 2) generated from IFNγ-/- mice or with anti-IFNγ mAb (Group 3). Data from these groups is pooled from 2 independently performed experiments. B-J. For CRS characterization, CRS associated cytokines IL-6, TNFα and IFNγ were analyzed in Group 1 (B, E, H), Group 2 (C, F, I) and Group 3 (D, G, J) respectively. Pre and post cytokine levels are paired values taken from each mouse alive at week 4 per cohort (Group 1 N=20, Group 2 N=13, and Group 3 N=12). K-M Flow cytometric analysis of macrophages was done using a subset of mice alive at week 4 from Groups 1-3 (Group 1 N=11, Group 2 N=10, Group 3 N=6). The frequency of (K) total macrophages (Live+CD45+CD11b+F4/80+), (L) iNOS+M1-like macrophages (Live+CD45+CD11b+F4/80+ CD11c-CD86+MHCII+iNOS+) and (M) Arginase1+ M2-like macrophages (Live+CD45+CD11b+F4/80 CD11c-+CD206+Arginase1+) are represented as a percent (%) of CD45+ cells. N-S. For characterizing neutrophils, cytokines G-CSF (N, P, R) and IL-17A (O, Q, S) were evaluated in Groups 1-3, respectively. Paired cytokine values for IL-17A and G-CSF are from 1 experiment (Group 1, N=8, Group 2 N=6, Group 3 N=5). T-V Flow cytometry of neutrophil apoptosis and proliferation were characterized in a subset of mice alive at week 4 for Group 1 (N=10 for apoptosis and N=6 for proliferation), Group 2 (N=6 for apoptosis and N=3 for proliferation) and Group 3 (N=3 for apoptosis and proliferation). Bar plots represent percentage (%) of (T) neutrophil apoptosis (Annexin V+ cells as a frequency of purified neutrophils) and (U) neutrophil proliferation (BrDU+ cells as a frequency of purified live neutrophils). (V) % Ly6G+CXCR2+ mature neutrophils (gated on Live+Lineage-CD11b+Gr1+ckit-CXCR4-BMMCs) was measured in a subset of mice alive at week 4 (Group 1, N=8, Group 2 N=4, Group 3 N=6). (W) Bar plots for % GMP cells are quantified in a subset of mice alive at week 4 by gating on Live+ckit+Sca1-FcγR+CD34+ BMMCs in groups 1-3 (Group 1 N=6, Group 2 N=8, Group 3 N=3). Error bars represent SEM. P values *P < .05, **P < .01, and ***P < .001 were considered significant. P values for cytokine bar plots B-J and N-S are derived from paired t test. P values for plots K-M and T-W are derived from Tukey’s multiple comparison test (using One way ANOVA).

CAR-T treated KO mice (Group 1) reproduced elevation of IL-6 and TNFα, but these cytokines were reduced in IFNγ-/- CAR-T and anti-IFNγ mAb groups (Fig. 4 B-G). The increase in IFNγ post CAR-T was observed again after IFNγ mitigation likely to the TMDD effect or secretion by other immune cells (Fig. 4 H-J). Analysis of bone marrow derived macrophages corroborated the significance of IFNγ blockade to alleviate CRS. CAR-T administration in Group 1 increased the total macrophage population (F4/80+CD11b+) (p=0.032) (Fig. 4 K). These bone marrow derived macrophages (from Group 1) were distinctly polarized towards the M1-like phenotype (CD86+MHCII+iNOS+) (p=0.004), where iNOS expression is associated with CRS (Fig. 4 L), as compared to the M2-like phenotype (CD206+Arginase1+) (Fig. 4 M). There was a significant reduction in M1-like macrophages with either mode of IFNγ-blockade (Fig. 4 L).

The elevation of cytokines in Group 1 coincided with a significant decline in cytokines associated with neutrophil survival, G-CSF (p=0.019) (Fig. 4 N) and IL-17A (p=0.030) (Fig. 4 O). IL-17A stimulates fibroblasts that release G-CSF to activate the transcription factors such as Gfi1 and CEBPα crucial for granulopoiesis [37–42]. IFNγ inhibition supported a recovery of G-CSF and IL-17A levels to baseline in group 2 (Fig. 4 P, Q) and group 3 (Fig. 4 R, S). Endpoint analysis revealed a significant reduction in the percentage of bone marrow derived apoptotic neutrophils (Fig. 4 T) with IFNγ-blockade in group 2 and group 3. This corresponded with a significant increase in neutrophil proliferation (Fig. 4 U) in group 2 (p=0.048) and group 3 (p=0.012). The inhibition of mature neutrophils (Fig. 4 V) in group 1 showed a significant recovery with IFNγ mitigation (Group 2, p=0.040, Group 3, p=0.034). The increase in neutrophil apoptosis and poor neutrophil recovery in CAR-T treated KO mice prompted an investigation into the impact of cytokines on granulocyte-monocyte progenitors (GMP) or granulocyte progenitors (GP). GMP and GP cells were FACS sorted using lineage depleted bone marrow cells of KO mice 4 weeks after treatment with CAR-T cells or IFNγ-/- CAR-T cells. The sorted GMP/GP cells were cultured ex vivo and their colony formation potential was measured (Fig. S4 A). CAR-T treated KO mice had significantly dampened GMP/GP colony formation not observed in WT mice or KO mice treated with IFNγ-/- CAR-T cells (Fig. S4 B-D). Flow cytometric analysis of GMPs shown as percent CD34+FcγR+ cells in bone marrow confirmed a significant drop in GMP population from CAR-T treated KO mice (Fig. 4 W), while IFNγ-blockade restored neutrophil progenitors to pre-CAR-T levels.

### IFNγ Modulation of the Th1/Th17 Axis Drives CRS and Neutropenia

To delineate the influence of IFNγ on Th1/Th17 balance and neutrophil homeostasis (Fig. 5A) naïve CD4+ CD62Lhi CD44low cells were isolated from C57BL/6 (B6) mice and confirmed to have no expression of the Th1 cytokine IFNγ or the Th17 cytokine IL-17A (Fig. S5 A-B). Isolated naïve CD4+ T cells were cultured with Th1 or Th17 inducing cytokines followed by co-culture with CD1928ζ CAR-T cells or IFNγ- /- CD1928ζ CAR-T cells (Fig. S5 C). Differentiation of Th0 to Th17 was 38.1% when cultured without CAR T cells (Fig. S5 D). When these cells were co-cultured with CD1928ζ CAR-T differentiation to Th17 dropped to 1.9%. Co-culture with IFNγ-/- CAR-T cells supported 17.3% Th0 cells to differentiate into Th17 (Fig. S5 D). Without CAR T cells 85% Th0 cells differentiated into Th1, which dropped to 59.75% in presence of CD1928ζ CAR-T cells and 8.29% in presence of IFNγ-/- CAR-T.

**Figure 5:**
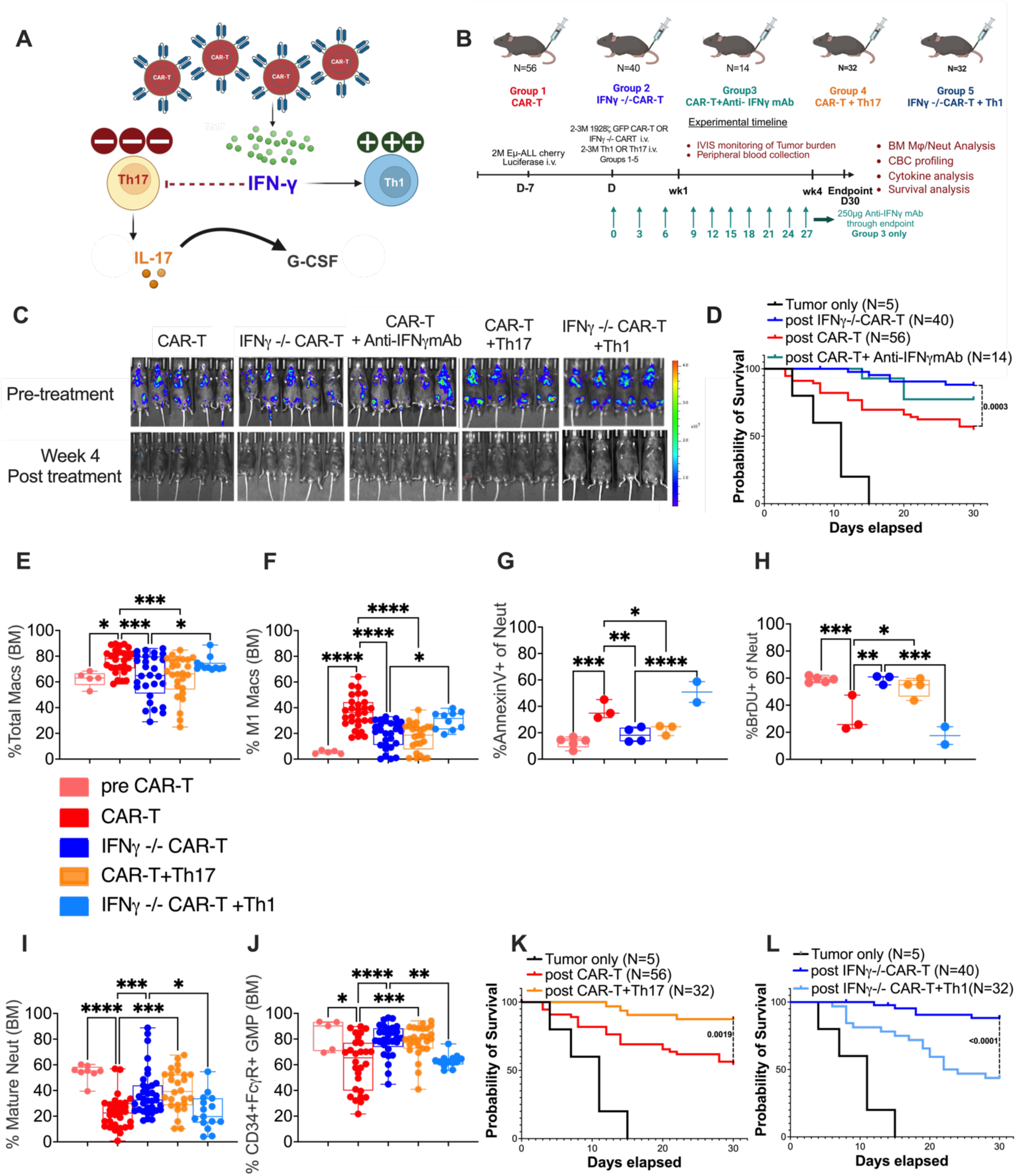
The impact of IFNγ blockade on the Th1/Th17 axis post CAR-T administration. A. Graphical representation demonstrating the proposed mechanism for the role of IFNγ in regulating Th1/ Th17 axis during CRS and neutropenia. B. KO mice bearing Luciferase expressing Eμ-ALL tumors were administered with CAR-T (Group1), IFNγ-/- CAR-T (Group 2), CAR-T treated with anti-IFNγ mAb (Group 3), CAR-T + Th17 cells (Group 4) and IFNγ-/- CAR-T with Th1 cells (Group 5). A control group included mice injected with Tumor only. Mice were imaged pre-treatment and at week 4 (endpoint) followed by analysis of CBC, neutropenia and CRS. Data in groups 1, 2, 4, and 5 are pooled from 3 independently performed experiments. Group 3 and Tumor only are from 1 of the pooled experiments so statistical analysis for these groups are not included. C. Representative IVIS images showing tumor burden in respective groups pre-treatment and at endpoint. D. Difference in Kaplan-Meier overall survival of mice between Group 1, Group 2 and Group 3. % Total macrophages (Live+CD45+CD11b+F4/80+) (E) and % iNOS+M1-like macrophages (Live+CD45+CD11b+F4/80+CD11c-CD86+MHCII+iNOS+) (F) were compared. G. % Annexin V+ apoptotic cells as a frequency of neutrophils (kit-based purification), (H) % BrDU+ cells as a frequency of live neutrophils (kit-based purification), and (I) % Ly6G+CXCR2+ mature neutrophils (gated on Live+Lineage-CD11b+Gr1+ckit-CXCR4-BMMCs) were evaluated for assessing neutrophil recovery. J. % GMP cells were determined by gating on Live+ckit+Sca1-FcγR+CD34+ BMMCs. K-L. Difference in Kaplan-Meier overall survival between Groups 1 and 4 (K) and Groups 2 and 5 (L) was assessed. Mice used in Groups 1, 2, and Tumor only are the same in D, K and L. Mice alive at endpoint were used to determine macrophages (Group 1 N=29, Group 2 N= 28, Group 4 N=26 and Group 5 N=10), mature neutrophils (Group 1 N=31, Group 2 N= 34, Group 4 N=26 and Group 5 N=14) and GMPs (Group 1 N=28, Group 2 N= 29, Group 4 N=26 and Group 5 N=13) in E,F,I, J. Annexin V and BrDU staining was done using a subset of mice from Group 1 (N=3 for Annexin V and BrDU), Group 2 (N=4 for Annexin V and N=3 for BrDU), Group 4 (N=3 for Annexin V and N=4 for BrDU), and Group 5 (N=2 for Annexin V and BrDU) in G and H. Error bars represent SEM. P values *P < .05, **P < .01, and ***P < .001 were considered significant. P values for Kaplan-Meier survival curves D,K,L were generated using Log-rank (Mantel-Cox) test. P values for box plots E-J were generated using Tukey’s multiple comparison test (using One way ANOVA).

To further interrogate the role of the Th1/Th17 axis on CAR T function IL-2Rα-/- mice were inoculated with syngeneic Eμ-ALL that expressed luciferase (Fig. 5 B). A week later mice with established tumors were divided into 5 groups. In addition to Group 1 (CAR-T), Group 2 (IFNγ-/- CAR-T), and Group 3 (CAR-T +anti-IFNγ mAb); Group 4 included CAR-T and Th17 cells that were adoptively transferred at a 1:1 ratio. Group 5 mice were administered IFNγ-/- CAR-T followed by an equal number of Th1 cells. As reported by Bailey et. al. [20], we confirm that IFNγ blockade does not alter the anti-tumor efficacy of CD1928ζ CAR-T cells (Fig. 5 C). Moreover, IFNγ-blockade improved the survival of mice post CAR-T infusion (p=0.0003 for Group 1 vs Group 2, and p=0.136 for Group 1 vs Group 3) (Fig. 5 D). We also noted total and M1-like macrophages are significantly reduced by IFNγ blockade (Fig. 5 E, F). There were no significant changes in the frequency of M2-like macrophages in Groups 1, 2, and 3 (Fig. S6 A). However, CAR-T cells induced iNOS in M2-like macrophages that was reduced with IFNγ blockade (Group 2) (Fig. S6 B). These observations reproduce our recent report showing CAR-T induced their own suppression by upregulating iNOS in M2-like macrophages, which could be reversed when blocking IFNγ (*<u>Lee et. al.,</u> <u>Preprint citation to be added</u>*). We also noted increases in apoptotic neutrophils and decreases in neutrophil proliferation, maturation and GMP frequency, which could all be reversed by IFNγ attenuation (Fig. 5 G-J).

We hypothesized that Th0 to Th1 differentiation caused by IFNγ released during CAR-T treatment results in an inadequate reserve of Th0 cells for Th17 differentiation. This is confirmed by addition of Th17 cells to CAR-T treated mice (Group 4), which displayed reduced neutrophil apoptosis, enhanced neutrophil proliferation and maturation (Fig. 5 G-I). Th17 adoptive transfer also led to increase in GMPs (Group 1 vs Group 4: p=0.0002) (Fig. 5 J). This improvement of BM neutrophil homeostasis due to Th17 adoptive transfer increased neutrophil concentration in peripheral blood compared to CAR-T alone (Fig. S6 C, E). Moreover, Th17 cells reduced total and M1-like macrophages (Fig. 5 E, F) and signficantly improved the survival of CAR-T treated mice (Fig. 5 K). We hypothesized that IFNγ released by CAR-T cells increases Th1 differentiation and M1-like macrophages, which was validated through Group 5 showing adoptive transfer of Th1 cells reversed the benefit of IFNγ blockade on toxicities by triggering polarization of M1-like macrophages (Group 2 vs Group 5: p=0.049) (Fig. 5 F). Th1 adoptive transfer also led to an increase in apoptotic neutrophils and a decrease in proliferative and mature neutrophils compared to the IFNγ-/- CAR-T group (Fig. 5 G – I). IFNγ-/- CAR-T treatment showed no difference in peripheral blood neutrophils (Fig. S6 D), whereas adoptive transfer of Th1 resulted in a significant drop in neutrophil concentration (Fig. S6 F). Re-establishment of CRS and neutropenia in IFNγ-/- CAR-T treated mice infused with Th1 cells resulted in poor survival (p<0.0001) (Fig. 5 L).

### Single cell Characterization of Bone Marrow Neutrophils and Macrophages in IL2Rα-/- Mice During CRS and Neutropenia

We performed single cell sequencing to compare differences in Neutrophils and Macrophages from BMMCs isolated at week 4 from Group 1 (CAR-T, n=2), Group 2 (IFNγ-/- CAR-T, n=2), Group 4 (CAR-T+Th17, n=2) and Group 5 (IFNγ-/- CAR-T +Th1, n=2). We used uniform manifold approximation and projection (UMAP) [43] to represent every cell in a two-dimensional embedding, which facilitates cell type identification based on shared gene expression markers in clusters of cells (informed by data from the Immunological Genome Project, ImmGen [44]). UMAP plots for each sample showcase different cell types, their respective numbers, and their molecular heterogeneity with nearly 70% of BMMCs in IL2Rα-/- mice being either neutrophils or macrophages/monocytes (Fig. 6 A, S7 A). Frequency of neutrophils are improved from Group 1 (16.9%) to Group 2 (64%) owing to IFNγ attenuation (Fig. 6 B). Similarly, Th17 administration in Group 4 increased neutrophils (63.2%) (Fig. S7 B). Conversely, Th1 administration in Group 5 reduced neutrophils (50.6%) (Fig. S7 B) compared to Group 2 (IFNγ-/- CAR-T). We also note macrophages in Group 1 (41.45%) were reduced by IFNγ mitigation in Groups 2 (15%) and 4 (17.9%) (Fig. S7 B). Th1 administration in Group 5 (19.2%) yielded a small increase compared to IFNγ-/- CAR-T Fig. S7 B).

**Figure 6:**
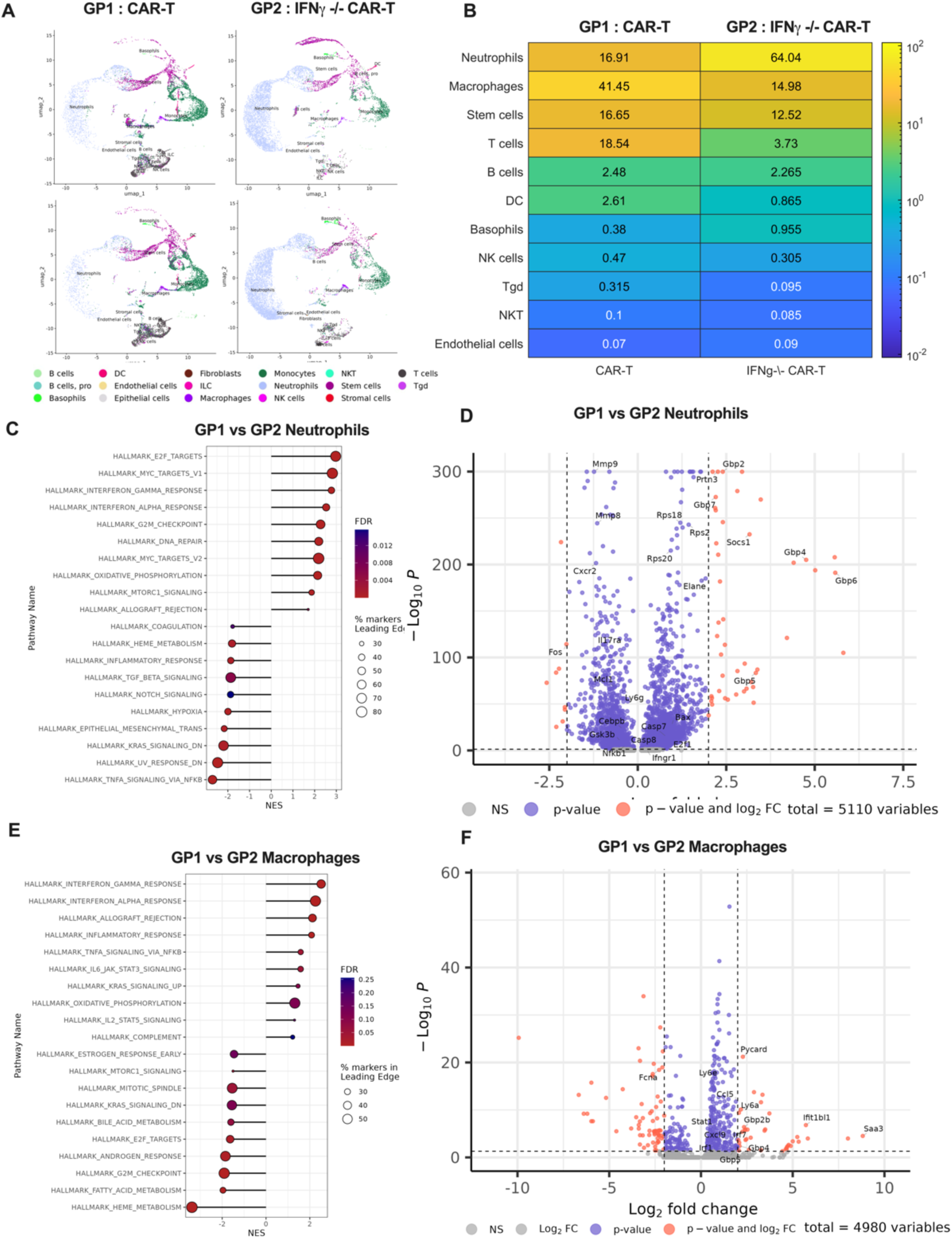
Utilizing Single cell sequencing to identify phenotypic differences in bone marrow neutrophils and macrophages from CAR-T and IFNγ-/- CAR-T treated mice. A. UMAPs representing cell types present in bone marrow samples of CAR-T treated (Group 1 or GP 1) and IFNγ-/- CAR-T treated (Group 2 or GP 2) samples. Each plot represents an individual mouse with 2 mice per Group. B. Heat map displaying the frequency of immune and non-immune cells cells in GP 1 and GP 2. Computed values are an average of 2 mice per Group. C. Gene set enrichment analysis (GSEA) showing the various Pathways from Hallmark dataset that were enriched in the comparison of Neutrophils between GP 1 and GP 2. D. Volcano plots highlighting the most differentially expressed genes in the Neutrophils from GP 1 and GP 2. E. GSEA showing various Pathways from Hallmark dataset that were enriched in the comparison of Macrophages between GP 1 and 2. F. Volcano plots highlighting the most differentially expressed genes in the Macrophages from GP 1 and 2.

Gene set enrichment analysis (GSEA) [43] of cancer hallmarks pathways in neutrophils showed upregulation of IFNγ response genes in Group 1 compared to Group 2 (Fig. 6 C), which is confirmed by higher expression of IFNγ-inducible guanylate binding proteins (Gbp2, Gbp4, Gbp6, and Gbp7) [45] (Fig. 6 D). This correlated with a downregulation of transcription factors involved in neutrophil activation and emergency granulopoeisis (Nfkb1, Gsk3b and Cebpb) [46–48]. Furthermore, overexpression of Prtn3, previously observed in apoptotic neutrophils [49], in addition to degradation of c-Fos [50] are indicative of increased apoptosis being triggered in Group 1 neutrophils. An upregulation of cell cycle gene sets involving E2F is attributed to overexpression of E2f1, and MYC due to overexpression of Casp8 and Bax (Fig. 6 C - D) that drive MYC-induced apoptosis [51–53]. Upregulation of E2f1 is associated with overexpression of Casp7 and underexpression of Mcl1 in neutrophils of Group 1 compared to Group 2, which support E2f1’s role in apoptosis [54–57]. However, Group 2 neutrophils are enriched for heme metabolism, hypoxia and TNFα signaling via NFKB (Fig. 6 C), which promote neutrophil survival [58–60]. We also note that Group 1 macrophages are enriched for IFNγ and IFNα response pathways compared to Group 2 (Fig. 6 E), which is attributed to upregulation of Ccl5, Stat1 and Fgl2 genes involved in M1-like macrophage polarization via the Cxcl9/Cxl10 axis (Fig. 6 F) [61–63] (*<u>Lee et. al., Preprint citation to be added</u>*). We observed over expression of interferon regulatory factors, IRF1 (induced by IFNγ) and IRF7 (induced by IFNα), which in turn aid in the production of iNOS [64–66]. Collectively, these analyses offer insight into putative molecular mechanisms involved in excess M1-like macrophage polarization, and subsequent activation of neutrophil apoptosis during CRS and neutropenia co-occurence. These molecular signatures are also seen in Group 1 (CAR-T) vs Group 4 (Th17 adoptive transfer) and Group 2 (IFNγ -/- CAR-T) vs Group 5 (Th1 adoptive transfer) comparisons (Fig. S8 A-H).

### Association of Th1/Th17 Cytokine Levels in Human Serum with Co-occurrence of CRS and Neutropenia in CAR-T Treated Patients

We investigated if Th1/Th17 balance is associated with CAR T cell toxicities in 27 patients treated at the Roswell Park Comprehensive Cancer Center (Supplementary Table S1). These patients were graded for CRS based on ASTCT guidelines [67]. We utilized a neutropenia classification by Rejeski et al[68] (Supplementary Table S2) that categorizes neutropenia based on rate of recovery (Quick, Intermittant and Aplastic) and another based on severity (Protracted, Profound, Prolonged) per ASCO/IDSA consensus guidelines [69]. To determine the impact of Th1/Th17 we split the 27 patients into three groups: no co-occurrence of CRS-neutropenia, low-grade CRS-neutropenia, and high-grade CRS-neutropenia. Low-grade CRS-neutropenia included patients with co-occurrence of neutropenia and mild or grade 1 CRS. High-grade CRS-neutropenia included patients with grade 2 or higher grade CRS, and prolonged or aplastic neutropenia (severe/high-grade neutropenia).We considered grade 2 CRS as high grade since it requires clinical intervention [13]. Five patients experienced high-grade CRS-neutropenia, six patients experienced low-grade CRS-neutropenia and the remaining sixteen patients experienced no co-occurrence of CRS-neutropenia. We measured the ratio of IFNγ (induced by Th1 cells) to IL-17A (induced by Th17 cells) cytokine levels at pre-lymphodepletion and peak-post-infusion. A significant increase in IFNγ-to-IL-17A ratio was noted between high-grade CRS-neutropenia and no co-occurrence (p=0.023) as well as low-grade CRS-neutropenia (p=0.018) groups at peak-post-infusion (Fig. 7 A). Consistent with our in vivo studies we observe co-occurrence of CRS-neutropenia is associated with Th1/Th17 balance.

**Figure 7:**
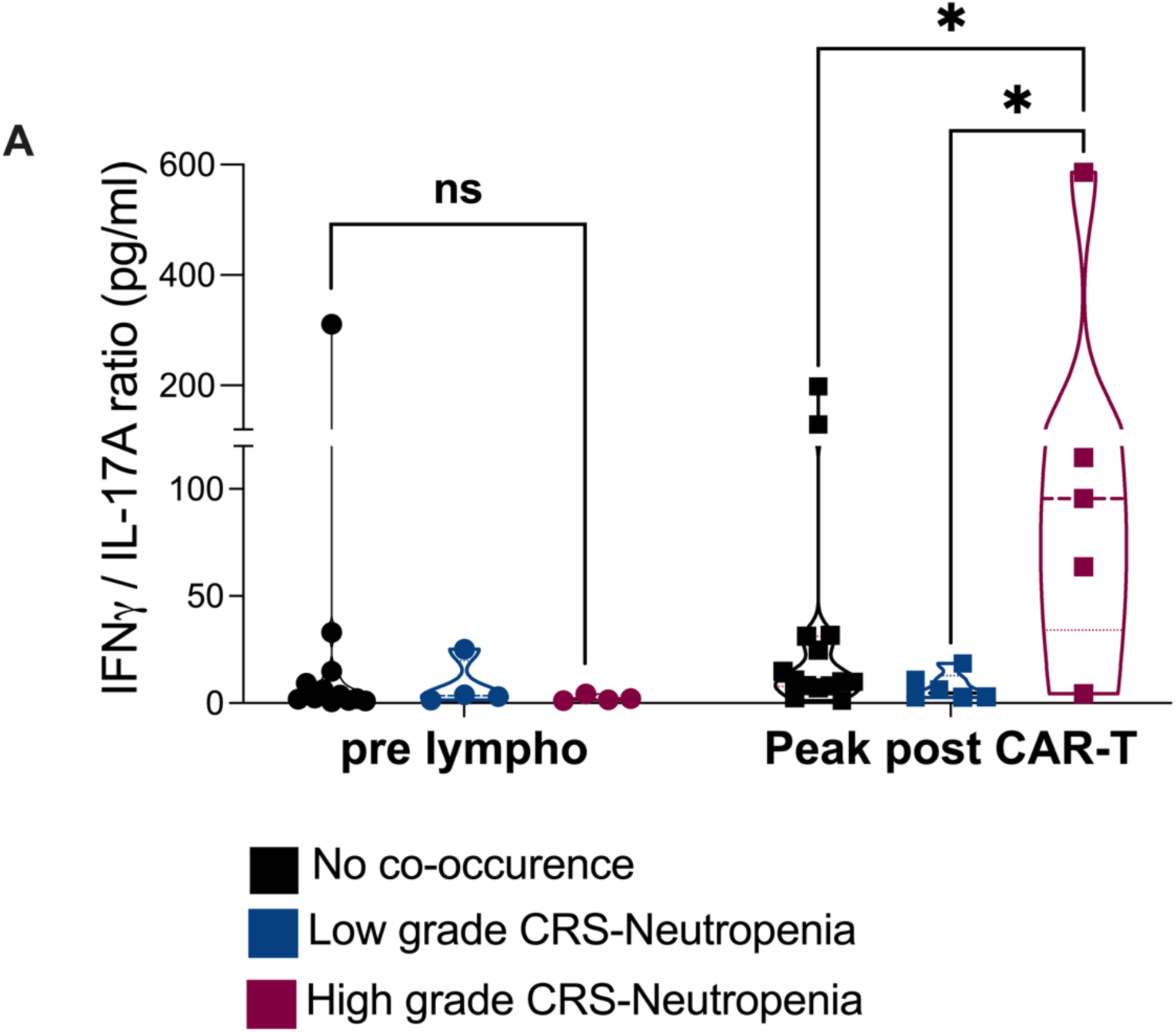
Determining the association of Th1/Th17 balance with CRS-Neutropenia co-occurrence in patients. A. Ratios of IFNγ to IL-17A (IFNγ/IL-17A) were calculated for 27 patients with hematological malignancies. Individual levels of IFNγ as well as IL-17A at prelymphodepletion (pre lympho : 6 days prior to CAR-T infusion) and peak post CAR-T infusion (defined as the highest cytokine value from post-infusion serum collected on Days 1, 7 and 11) were used to generate the ratio of IFNγ to IL-17A. Violin plots represent three different groups where the plots in black represent patients that did not develop co-occurrence of CRS and neutropenia (N=16), blue violin plots represent patients with a mild case of CRS and neutropenia co-occurrence (N=6), and red violin plots represent patients with a severe case of CRS and neutropenia co-occurrence (N=5). *P < .05, **P < .01, and ***P < .001 were considered significant and derived from unpaired t tests.

## Discussion

The advent of CAR-T therapy has greatly improved outcomes for patients with hematological malignancies, however a subset of patients continue to experience toxicities such as CRS, ICANS, and ICAHT [1]. Rejeski et al show that 51% of non-relapse patient mortality (NRM) is due to infections attributed to cytopenias [12]. There are currently no pre-clinical in-vivo models that recapitulate cytopenias and while mouse models for CRS exist [32, 33] they are not immune competent and not ideal to study cytopenias. We employed a IL-2Rα-/- *in vivo* model to recapitulate CRS in Eμ-ALL tumor-bearing mice and observed a significant increase in pro-inflammatory cytokines and death from CRS. Non-tumor-bearing KO mice treated with CAR-T also exhibited CRS, which either anti-IL-6R or anti-IFNγ mAb alleviated. The predominant manifestation of ICAHT is prolonged neutropenia characterized by a hypocellular bone marrow that is non-responsive to growth factors and is without effective treatment [6]. ICAHT is posited to be related to cytokines since clinically CRS and neutropenia co-occur, and ICAHT after CAR T cell therapy is categorically different than the neutropenia observed from chemotherapy alone. [6, 7, 9]. Similar to patients CAR-T treatment in the IL-2Rα-/- model resulted in prolonged reduction of neutrophils, while lymphodepletion-induced neutropenia had rapid recovery. Prolonged neutrophil reduction associated with CAR-T treatment was shown to coincide with a dramatic disruption of bone marrow neutrophil homeostasis. This was accompanied by an increase in pro-inflammatory M1-like macrophages.

We investigated the role of IFNγ-blockade on CAR-T treated KO mice in two ways: by combining CAR-T with anti-IFNγ mAb resulting in a systemic inhibition of IFNγ and by employing IFNγ-/- CAR-T for a more targeted effect. Using either form of IFNγ-blockade we noted an alleviation of CRS and neutropenia, thereby implicating excess IFNγ produced by rapidly expanding CAR-T cells as a key mediator of these toxicities. Most notably, we observed no significant differences in CAR-T efficacy with IFNγ-blockade, and between both systemic (using anti-IFNγ mAb) and targeted (IFNγ-/- CAR-T) blockade of IFNγ, which is consistent with findings from a recent study involving hematological malignancies [20]. CAR-T associated CRS is characterized by an increase in cytokines IFNγ and TNFα, while neutropenia is characterized by a decrease in IL-17A and G-CSF. Th17 cells produce IL-17, which regulates G-CSF [70] responsible for granulopoiesis, while Th1 cells produce IFNγ [71] that supports pro-inflammatory M1-like macrophages [72] leading to CRS. Both Th1 and Th17 cells originate from naïve CD4+ T cells, which differentiate into Th1 cells in the presence of IL-12 and IFNγ [<u>73</u>], while Th17 differentiation is supported by TGFβ and IL-6 [74]. Tregs play a crucial role in balancing the Th1-Th17 axis by producing TGFβ aiding in Th17 differentiation [75] and IL-10 that suppresses antigen presenting cells (APC) from releasing IL-12 [76]. IL-2Rα-/- in C57BL/6 mice leads to Treg dysfunction resulting in breakdown of such negative feedback loops. In the absence of Tregs, IFNγ released by CD8+ T cells induce APCs to produce IL-12 leading to Th1 differentiation, more IFNγ secretion, and a positive feedback loop [77]. Systemic and targeted IFNγ-inhibition prevent uncontrolled Th1 differentiation and may temporarily re-instate homeostatic immune function and cytokine balance.

Using single-cell RNA sequencing we characterized bone marrow neutrophils and macrophages, which revealed upregulation of apoptotic signaling in neutrophils and overexpression of markers associated with M1-like polarization in macrophages in IL2Rα-/- mice treated with CAR-T. The role of Th1/Th17 balance driving CRS and neutropenia is verified with single-cell RNA sequencing where similar molecular characteristics were observed after adoptive transfer of Th1 cells with IFNγ-/- CAR-T compared to CAR T cells alone and IFNγ-/- CAR-T treatment compared to adoptive transfer of Th17 cells. Moreover, the role of Th1/Th17 imbalance in the context of co-occurrence of CRS-neutropenia is shown using patient cytokines. Peripheral blood samples of patients treated with CAR-T show an increase in IFNγ-to-IL-17A ratio between high-grade CRS-neutropenia and no co-occurrence of CRS-neutropenia. This is also confirmed by a recent study that shows a similar decrease in IL-17A to be associated with CRS and ICANS [78].

Our work identifies the downstream mediators of IFNγ responsible for CRS and neutropenia that lead to adverse patient outcomes.The use of IFNγ blockade to clinically abrogate cytokine-mediated toxicities after CAR-T cell therapy is being studied in a clinical trial (NCT06550141). While the trial is administering anti-IFNγ mAb prior to CAR-T, our work shows the benefit of IFNγ blockade in alleviation of CRS and ctyopenias after CAR-T administration. Currently, there is insufficient evidence to understand the impact of IFNγ blockade on the tumor-immune microenvironment prior to CAR-T therapy. IFNγ has a pleiotropic role on anti-tumor immunity so we speculate that its blockade during CAR-T therapy could have different effects on toxicities and efficacy based on when and how it is blocked. For example, we recently demonstrate that IFNγ secreted by CAR-T cells induce iNOS in M2-like macrophages that lead to CAR-T suppression and IFNγ blockade improves CAR-T tumor killing (*<u>Lee et. al., Preprint citation to be added</u>*). On the other hand a study by Tang et. al. shows that lack of IFNγ maintains V-domain Ig suppressor of T-cell activation (VISTA) that inhibits tumor killing efficacy and persistence of CD19-CAR-T cells [79]. This pleitropic effect of IFNγ suggests that its blockade could have different effects from patient-to-patient based on features of the patients tumor, TME, and CAR-T cell product. This highlights the importance of understanding how each of these components are regulated by IFNγ when designing translational trials of IFNγ blockade to reduce toxicities and improve durable responses for patients treated with CAR-T cells.

## Materials & Methods

### Mice

All animal studies were conducted per protocols approved by the Institutional animal care and use committee (IACUC) of the Roswell Park and Moffitt Comprehensive Cancer Centers. IL2-Rα-/- mice (B6;129S4-Il2ratm1Dw/J, IMSR_JAX:002462) were obtained from JAX lab and bred in-house. Mice positive for neomycin allele and negative for wild type (WT) allele (-/-) were confirmed by genotyping as IL-2Rα-/- and used for *in vivo* studies. C57BL/6 (WT) mice were purchased by JAX laboratory and used as controls. Thy 1.1 (B6.PL-Thy1a/CyJ, IMSR_JAX:000406) were used as donors for the generation of CD19-28ζ CAR-T cells while IFNγ-/- mice (B6.129S7-IFNγtm1Ts/J, IMSR_JAX:002287) were used as donors for the generation of IFNγ-/- 19-28ζ CAR-T cells. Both male and female mice between 8 to 12 weeks old were used in experiments.

### Cell lines

Eμ−ALL cherry Luciferase expressing cells are a murine syngeneic B cell Acute lymphoblastic leukemia cell line generated in-house and maintained as described [19]. All cell culture reagents were purchased from Thermofisher. Mouse CAR-T cells, Naïve CD4+ T cells, Th1 and Th17 cells were cultured in RPMI-1640 supplemented with 10% FBS, 2 mmol/L L-glutamine, penicillin (100 U/mL), streptomycin (100 μg/mL) 2 mmol/L sodium pyruvate, 1 × nonessential amino acids, 10 mmol/L HEPES and 55 μmol/L β-mercaptoethanol. Universal Mycoplasma Detection Kit (ATCC) was used to test cell lines used in various experiments for mycoplasma.

### Genetic constructs and CAR-T cell generation

CD1928ζ CAR-T construct has been described [19, 80, 81]. The construct was used for retroviral production and T cell transduction per our established protocol [82]. Transduction efficiency was confirmed as a percent of total GFP+ or Cherry+ T cells (listed in graphical representations or legends in main and supplementary figures) as the CARs include a fluorescent reporter protein.

### In vivo studies

IL-2Rα-/- (KO) or C57BL/6 (WT) mice were injected intravenously (i.v.) with 1 million (M) to 2 M murine syngeneic Eμ−ALL cells (B cell Acute lymphoblastic leukemia) expressing luciferase. A week after tumor establishment mice were administered intraperitoneally (i.p.) with 200 to 300 mg/ kg of cyclophosphamide (Sigma-Aldrich) followed by infusion of 1 M to 3 M 19-28ζ CAR-T cells (also listed in graphical representations or legends for experimental design in main and supplementary figures as applicable). For Th1/Th17 adoptive transfer studies, Th1 or Th17 cells were injected at an equal ratio to CAR-T infusion.

For cytokine analysis, serum was collected from peripheral blood through the submandibular vein. Cytokines were measured using 25μL serum on a Luminex 100 system with a mouse luminex assay kit (R&D Systems) or Ella with a Simple Plex Assay Kit (Biotechne) according to manufacturers instructions.

For Histopathology mouse spleen and Lymph node tissues were collected at endpoint and stained using anti-B220 (CD45R) and anti-CD3 Antibodies. H & E staining was performed for spleen, lymph node, bone marrow and liver sections and examined using Leica light microscopy.

For analysis of peripheral blood, samples were subjected to ACK lysis buffer (GIBCO, Thermofisher scientific) for 15 minutes at room temperature followed by staining with anti-CD19 (BD Biosciences) to measure levels of B cells and CD3 as well as Thy1.1 marker to detect levels of Thy1.1+GFP+ CAR-T cells in peripheral blood (PBL).

For CRS mitigation using blockade of IL-6R and IFNγ, KO and WT mice were treated intraperitoneally with either 12.5 mg/kg (250-300 μg/mouse) of murine anti-IL-6R mAb (MP5-20F3; BioXcell) or 250 μg/mouse of murine anti-IFNγ mAb (Clone XMG1.2, Bioxcell) or Isotype control (Rat IgG1, clone HRPN, BioXcell) (N=21 for Figure 2) as described [83, 84]. Treatments were repeated twice weekly until endpoint.

For Treg adoptive transfer studies, murine CD4+CD25+Foxp3+ Tregs were isolated using a kit (STEMCELL) from the spleens of C57BL/6 mice per manufacturer instructions. Purified cells were resuspended in 1X PBS (100L per mouse) and injected into KO mice at a dose of 0.25 M Tregs or 1 M Tregs. Two million CAR-T cells were infused two days after transfer of Tregs.

For complete blood profiling to assess neutrophil concentration, 50 μL of peripheral blood was collected per mouse. The blood samples were measured using ProCyte Dx® Hematology Analyzer.

For BrDU staining, neutrophils were isolated from bone marrow of KO or WT mice pre and post infusion. Cells were counted and processed using the instructions provided by the kit manufacturer (BD Pharmingen). Briefly, cells stained with viability dye (Zombie NIR, Biolegend) were fixed and permeabilized using cytofix/cytoperm buffer for 30 minutes. The cells were then treated with DNAse (1mg/mL) and incubated for an hour at 37°C followed by staining with anti-BrDU (BD Phramingen) at 1:50 dilution.

For Annexin V staining, isolated neutrophils were counted and washed twice with 1X Dulbecco’s phosphate buffered saline (GIBCO) as per manufacturer instructions (BD Phramingen). The cells were resuspended in 1X Annexin V binding buffer and stained using 5 μL anti-Annexin V as well as 5 μL 7-AAD for 15 minutes at room temperature in the dark. Cathepsin treated cells were used as positive control. Stained samples were acquired using Cytek Aurora within 1 hour of preparation.

For neutrophil maturation analysis, 1M total BMMC per sample were stained for flow cytometry with Zombie NIR, anti-CD3, anti-Nk1.1, anti-ckit, anti-CD115, anti-Gr-1, anti-Ly6G, anti-CXCR2, anti-CXCR4 (Biolegend); anti-CD11b and anti-B220 (BD Biosciences). Mature Neutrophils were gated as Live+Lineage-(CD3-, B-, NK-) ckit-CD115-Gr1+CD11b+CXCR4-Ly6G+CXCR2+ based on a previous study [85].

For characterization of M1-like and M2-like bone marrow macrophages, 1M total bone marrow mononuclear cells (BMMC) per sample were stained for flow cytometry with Zombie NIR, anti-CD45, anti-F4/80, anti-CD64, anti-Ly6G, anti-Ly6C, anti-CD115, anti-MHCII, anti-iNOS, anti-Siglec-F (Biolegend); anti-CD11b, anti-CD11c and anti-CD86, anti-CD206 (BD Biosciences) and anti-Arginase1 (eBioscience). Based on a previous study [86] % Total macrophages were gated as CD45+CD11b+F4/80+, % M1-like bone marrow macrophages were gated as CD45+CD11b+F4/80+CD11c-MHCII+CD86+iNOS+ and % M2-like macrophages were gated as CD45+CD11b+F4/80+CD11c-CD206+Arginase1+ as a percent total of CD45+ cells.

### Colony formation assay (CFU) and Granulocyte monocyte progenitor (GMP) characterization

The CFU assay was performed by culturing FACS sorted GMP and GP cells in Methocult media (STEMCELL). BMMCs were first subjected to lineage depletion (Miltenyi Biotec) followed by FACS sorting of GMP (ckit+Sca1-CD16/32 (FcγR)+CD34+Ly6C-Flt3-CD115low) and GP (ckit+Sca1- CD16/32(FcγR)+CD34+Ly6C+Flt3-CD115low) populations using BD FACS Aria based on a previous publication[87]. Sorted cells were pooled using two (out of four) mice per group, to represent two replicates per group and resuspended at a 10X concentration of 0.2-0.3 M cells in IMDM + 2% FBS. The cells were seeded at a density of 20-30,000 cells in 1.1 mL Methocult per 35 mm plate prepared in duplicates per group. The duplicates for each group were then placed inside a 100mm plate with a 3^rd^ 35 mm plate filled with water. The plates were incubated for 7 days at 37°C and 5% CO2. Colonies formed were imaged using phase contrast microscope and counted at day 7 using an STEMvision Hematopoietic Colony Counter. For flow cytometric characterization of GMPs, BMMCs were stained using anti-CD3, anti-B220, anti-Nk1.1, anti-ckit, anti-CD16/32 (FcγR), anti-Ly6C, anti-CD115, anti-Sca1, anti-Flt3 (Biolegend) and anti-CD34 (BD Biosciences). % GMP cells were gated as ckit+Sca1-CD16/32 (FcγR)+CD34+based on the previous study [87].

### In-vitro Th1/Th17 differentiation assay

The differentiation protocol was set up as described [88]. Naïve CD4+ T cells isolated (Miltenyi biotec) were seeded in 24 well plates at a concentration of 0.5 M cells per mL. Th1 differentiation cytokines included IL-12 at 10ng/mL (R & D systems), IL-2 at 200IU/mL (Iovance Biotherapeutics) and anti-IL-4 at 1 μg/mL (Thermofisher). Th17 differentiation cytokines included IL-6 at 40ng/mL (R & D systems), TGFβ at 3ng/mL (R & D systems), anti-IL-4 at 1 μg/mL, anti-IL-2 and anti-IFNγ at 1 μg/mL (Thermofisher). The differentiated cells were cultured either alone or in presence of 0.5 M CAR-T cells or IFNγ-/- CAR-T cells respectively (in separate wells). On day 4, cells were incubated with PMA at 5ng/mL (Sigma-Aldrich) and Ionomycin at 1mg/mL (Sigma-Aldrich) for an hour at 37°C. This was followed by addition of 10mg/mL Brefeldin A (eBioscience) and incubation for 4 hours. Stimulated T cells were further stained for flow cytometric analysis using anti-CD4, anti-CD44, anti-CD62L, anti-IL17 and anti-IFNγ (Biolegend).

### scRNAseq methods

Single cells were isolated and then processed using the 10X Genomics Single Cell 3’ v3 kit according to the manufacturer’s instructions. Libraries were sequenced on the Illumina NovaSeq 6000 instrument. Raw sequencing data were processed using Cell Ranger (CR) (v6.0.0) pipeline to generate fastq files. Fastq files were aligned and quantified generating feature-barcode count matrices. Gene-barcode matrices containing Unique Molecular Identifier (UMI) counts are filtered using CR’s cell detection algorithm. Downstream analyses were performed mainly using Seurat (v5.0.0) (1) single-cell analysis R package. Eight single-cell RNA seq samples were individually read into a Seurat object to examine feature number, mitochondrial percentage and read count distributions within each sample. Cells with less than 500 features or greater than 7500 features or >15% mitochondrial content were filtered out. After normalizing and finding variables features from individual samples, all samples are then integrated using FindIntegrationAnchors function. The resulting data would serve for visualization purposes. Separately, samples are merged in a SingleR (v1.6.1) (2), a Bioconductor package, was then used to annotate individual immune cell types using ImmGen reference database (3) from celldex (v1.12.0) R package. Principal component analysis was used to detect and visualize highly variable genes. Using the RNA assay, data normalization and scaling were performed using Seurat’s SCTransform (4) function regressing against mitochondrial percentage. The data were then dimension-reduced via UMAP and clustered using Louvain algorithm for downstream visualization. Normalization, scaling, dimension-reduction, clustering and quantification is performed on specific cell populations (i.e. Neutrophils and Macrophages) to further detect subpopulation clusters based on their expression profiles. Differential gene expression analysis was performed using Seurat’s FindMarkers function on groups of interest using the Wilcoxon Rank Sum test.

Subsequent gene-set enrichment analyses were performed on comparisons of interest using GSEApreranked procedure implemented by the clusterProfiler package (v4.6.2) (5) using msigDB database (6) found in the msigdbr (v7.5.1) package. Mouse versions of Hallmark, C2 and C5-GO gene sets were used to perform pathway analyses. The avg_log2FC values (obtained from the FindMarkers results) are utilized as ranked values to perform pathway analyses using prerankedGSEA (7). Specific cell-level signatures scores are obtained using Seurat’s AddModuleScore function.

### Patient samples and data collection

Patient samples were collected following informed consent under an IRB-approved protocol I-57217 and used in accordance with the Declaration of Helsinki, International Ethical Guidelines for Biomedical Research Involving Human Subjects (CIOMS), Belmont Report, and U.S. Common Rule. We obtained serum samples from 27 patients who underwent CAR-T cell therapy at the Roswell Park Comprehensive Cancer Center. The patients received standard-of-care CAR-T products. Serum samples were collected at the following time points: pre-lymphodepletion (within 10 days prior to CAR-T infusion), Day 0 (pre-CAR-T), Day 1, Day 7, and Day 14 (post-CAR-T). Neutropenia classification followed criteria from a previous study [68] . Prolonged neutropenia was defined as an absolute neutrophil count (ANC) < 1000 cells/µL, measured at ≥ 21 days after CAR-T cell infusion. Aplastic neutropenia was characterized by severe neutropenia (ANC < 500 cells/µL) lasting ≥ 14 days.

### Statistical analysis

All Statistical evaluations were performed on GraphPad Prism 10 software (GraphPad Software, La Jolla, USA) with the aid of a Biostatistician, using two-tailed Mann–Whitney t-tests (Paired, Unpaired) or Tukey’s multiple comparison test using ANOVA analyses, in accordance with the type of data. All survival analyses were done using Log-rank (Mantel-Cox) test. p values < 0.05 were taken as significant values.

## Acknowledgements

We acknowledge the following shared resources at Roswell Park Comprehensive Cancer Center: Flow and Immune Analysis Shared Resource (FIASR), Lab animal shared resource (LARS), Genomics, Bioinformatics Shared Resource (BIOINF) and Experimental Tumor Model Resource. We would like to thank Dr. Renier Brentjen’s lab for providing us with Eμ-ALL cherry Luciferase expressing cells, Dr. Muhammad Junaid Tariq for patient sample collection and data procurement, Dr. Prashant Singh and Sarah McEvoy for performing single cell sequencing, Eduardo Cortes Gomez and Dr. Jianmin Wang for analysing single cell sequencing data. This study was funded by Hackensack meridian health-CDI, The Mark Foundation for Cancer Research, The Rustum Family Endowment for Translational Scientific Achievement and RO1 grant #NIH-5RO1AI155786-04.

## Author Contributions

**Payal Goala:** Writing – original draft, review and editing, Experiments – conceptualization, design and execution, Data – curation, visualization and analysis. **Yongliang Zhang**, **Nolan Beatty, Shannon McSain, Cooper Sailer, Justin C Boucher, Constanza Savid Frontera:** Experiment – execution, Writing-review and editing. **Muhammed Junaid Tariq, Showkat Hamid:** Patient sample collection and data procurement, Writing-review and editing. **Eduardo Cortes Gomez, Jianmin Wang:** Single-cell data analysis, Writing-review and editing. **Sae Bom Lee, Hiroshi Kotani:** Writing – review & editing. **Writing Michael D. Jain:** Conceptualization, Writing-review and editing. **Marco L. Davila:** Supervision, Funding acquisition, Conceptualization, Writing – original draft, review and editing.

## Disclosures

Zhang reports employment at Iovance Biotherapeutics. Michael D. Jain reports grants and personal fees from Kite/Gilead, personal fees from Myeloid Therapeutics, Novartis, and BMS, and grants from Incyte, Loxo@Lilly, FACCA, Mark Foundation, and Bankhead-Coley State of Florida outside the submitted work. Marco L. Davila reports fees from Kite, BMS, A2 Biotherapeutics, Poseida and stock and/or options in Adaptive Biotechnologies, Precision Biosciences, Adicet, and A2 Biotherapeutics. No disclosures were reported by the other authors.

