## Supplementary Figures for "The Th1/Th17 axis regulates chimeric antigen receptor (CAR) T cell therapy toxicities"

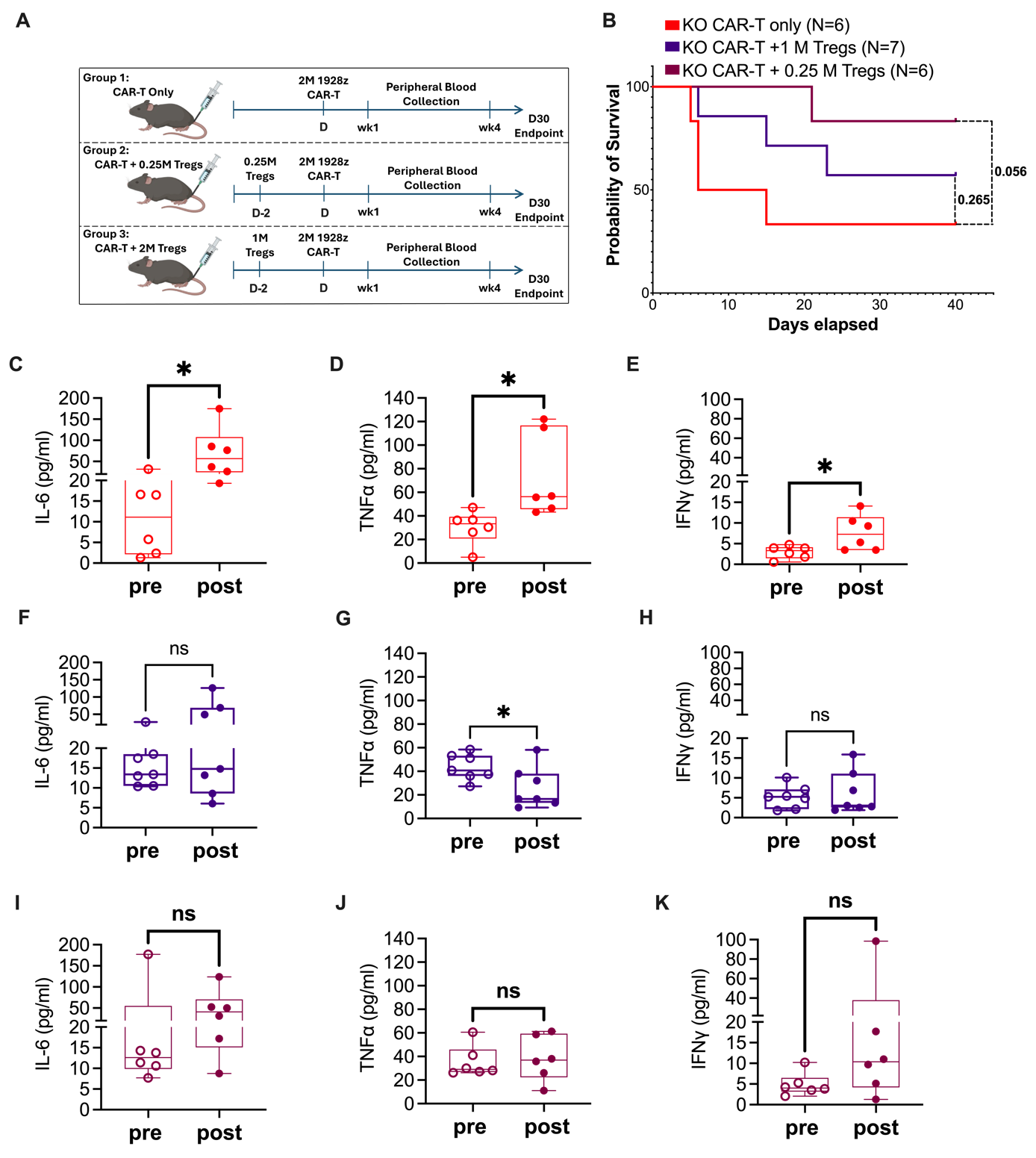
**Supplementary Figures**

**Figure S1:** A. Schematic diagram of 2M 1928ζ GFP CAR-T treated KO mice transferred in absence or presence of low (0.25 M) or high dose (1M) Tregs. Data represents a single experiment. B. Kaplan-Meier overall survival curve of CAR-T treated mice in presence or absence of Treg adoptive transfer. C-K. Cytokines IL6 (C, F, I), TNFα (D, G, J) and IFNγ (E, H, K) in CAR-T inoculated mice treated with 1 M Tregs comparing pre CAR-T vs week 4 levels. Pre and post cytokines are paired values taken from each mouse alive at week 4 or endpoint per mouse. Error bars represent SEM. P values *P < .05, **P < .01, and ***P < .001 were considered significant. P values for cytokine bar plots C-K were generated using paired t test. P values for Kaplan-Meier survival curve B was generated using Log-rank (Mantel-Cox) test.

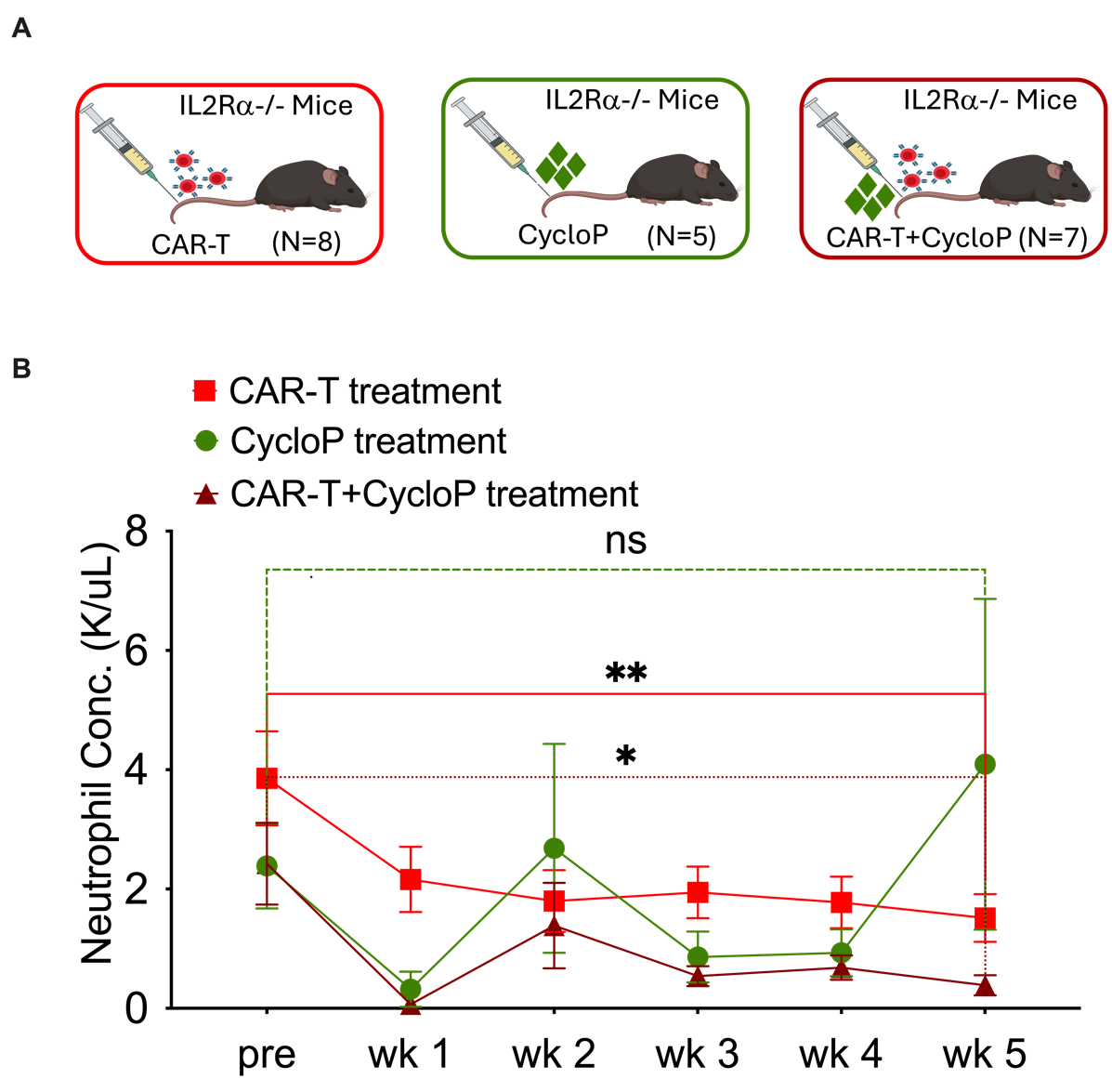
**Figure S2:** A. Schematic showing KO mice treated with either 2 M 1928ζ cherry CAR-T or 200 mg/kg cyclophosphamide (CycloP) alone or in combination. Differences in recovery rate of Neutrophils were assessed under these 3 different conditions in KO mice. KO mice infused with CAR-T alone are from one of the pooled experiments from Fig 3 A that included the two study arms of cyclophosphamide and CAR-T + cyclophosphamide. B. Comparison of neutrophil concentration in the respective groups. Data is pooled from 2 independently performed experiments. Error bars represent SEM. P values *P < .05, **P < .01, and ***P < .001 were considered significant. P values for line plot B was generated using paired t test.

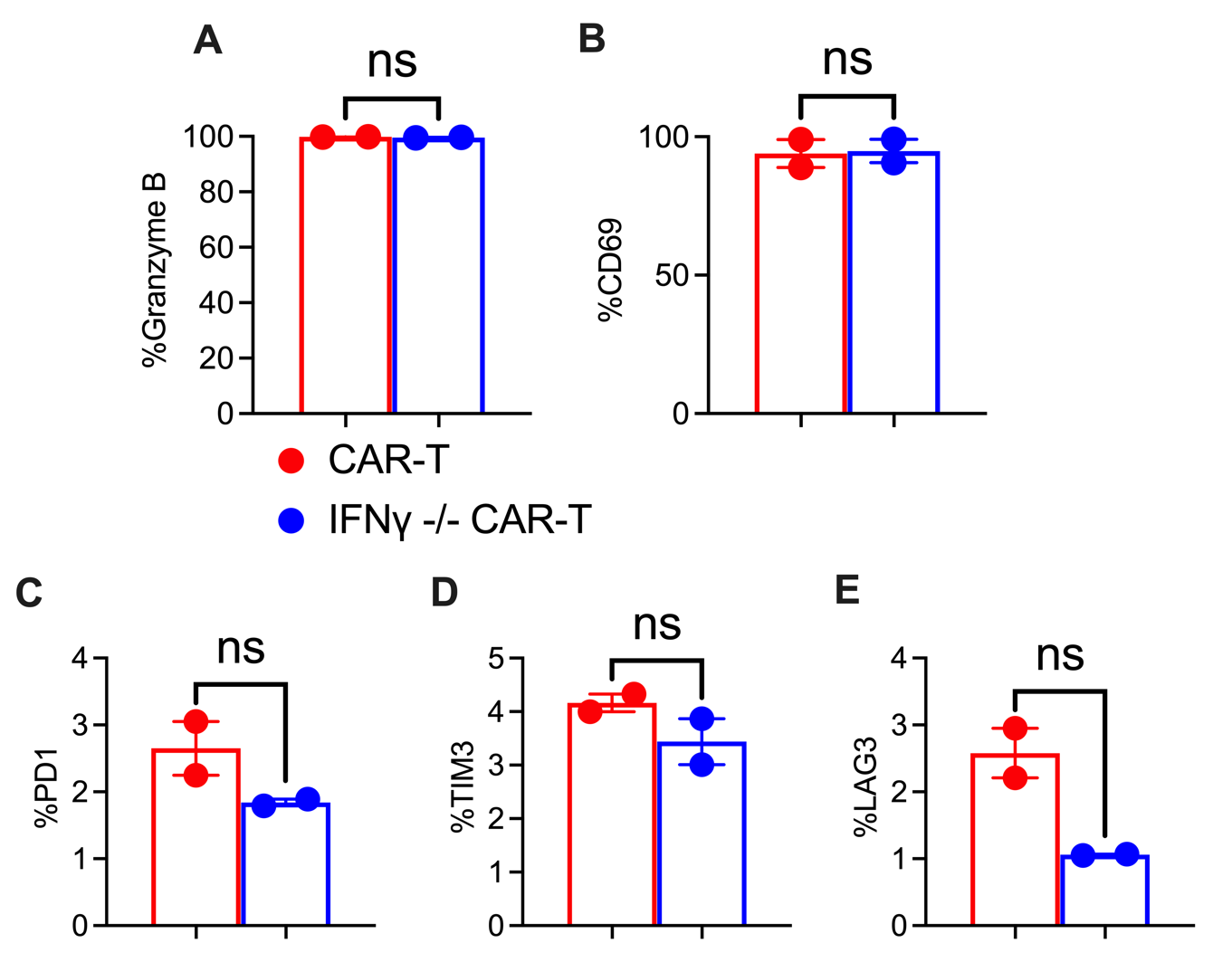
**Figure S3:** Day 5 transduced 1928ζ GFP CAR-T or IFNγ-/- CAR-T cells were debeaded and treated with Ionomycin for 1 hour, followed by Brefeldin A for 4 hours. Samples were stained intracellularly for activation markers Granzyme B (A), CD69 (B) as well as exhaustion markers such as PD1 (C), TIM3 (D) and LAG3 (E). Data are from one independent experiment (N=2). Error bars represent SEM. P values *P < .05, **P < .01, and ***P < .001 were considered significant. P values for bar plots A-E were generated using unpaired t test.

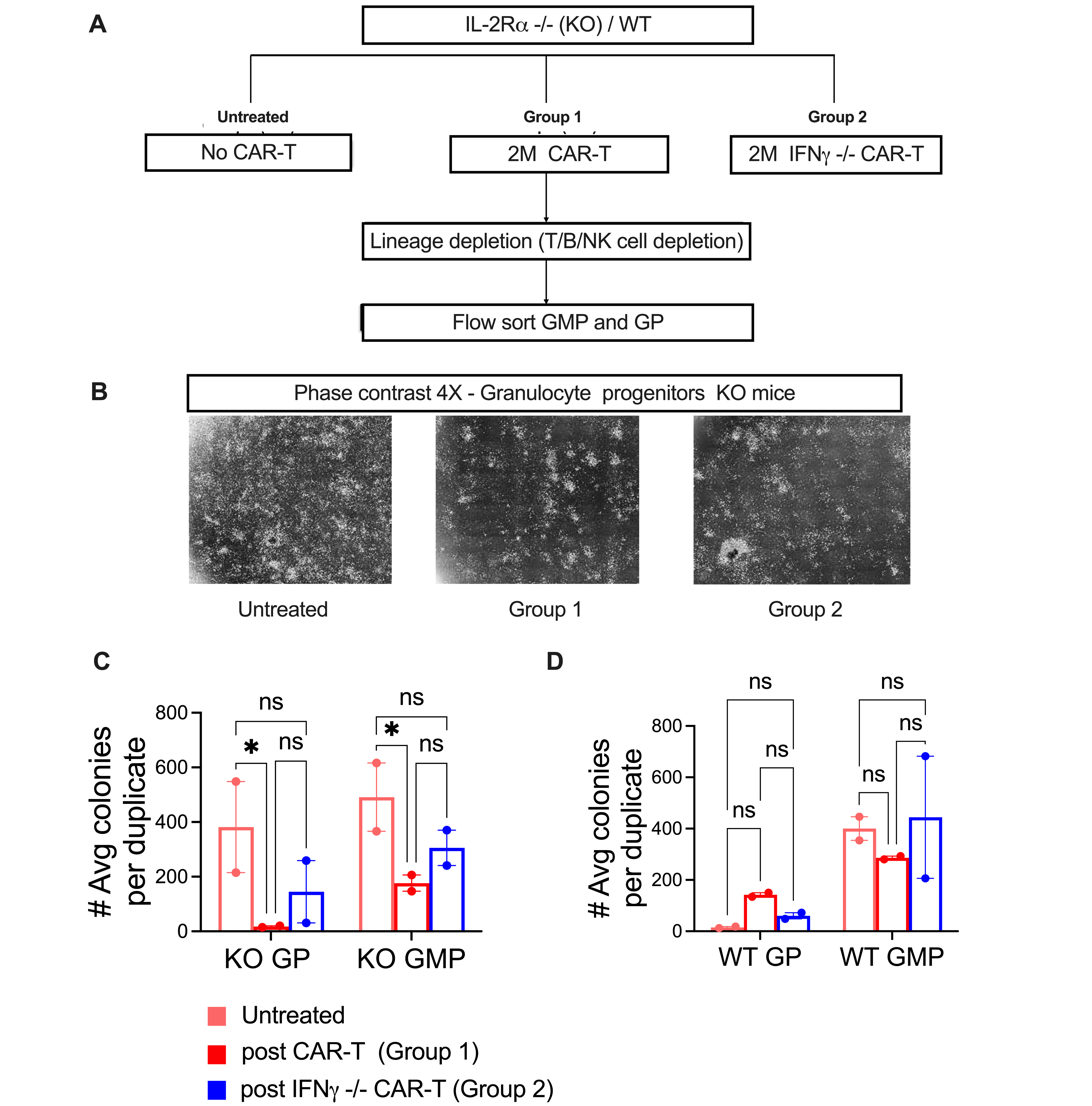
**Figure S4:** A. Diagrammatic representation of Granulocyte-monocyte-progenitor (GMP) and Granulocyte progenitor (GP) cell isolation for Methocult colony formation. Week 4 post 1928ζ GFP CAR-T (Group 1) or post IFNγ-/- CAR-T (Group 2) treated GMP and GP cells were enriched by Fluorescence-activated cell sorting (FACS) using BMMCs isolated from 4 KO and 4 WT mice per Group. Four untreated mice were used as control. BMMCs were depleted for lineage cells (T/B/NK cells) using kit-based lineage depletion. Lineage negative cell suspensions were FACS sorted to yield ckit+Sca1-FcγR+CD34+Ly6C-Flt3-CD115low GMP cells or ckit+Sca1-CD16/32(FcγR)+CD34+Ly6C+Flt3-CD115low GP cells as described (Methods). Data is represented from a single experiment (KO N=4 per Group, WT N=4 per Group). Sorted GMP cells were pooled from 2 of the 4 mice per group. The cells were cultured ex-vivo on Methocult media for 7 days. The same procedure was followed for GP cells. B. Representative images of a single plate per group displaying the GP colonies formed on Day 7 acquired using phase contrast microscope at 4X magnification. The number of GMP or GP colonies (plated separately) formed in each plate per group for KO (C) and WT (D), were counted on day 7 using an automated hematopoietic colony counter. Results were reported as average number of colonies formed from duplicates per group. Error bars represent SEM. P values *P < .05, **P < .01, and ***P < .001 were considered significant. P values for bar plots C and D were generated using Tukey’s multiple comparison test (using Two way ANOVA).

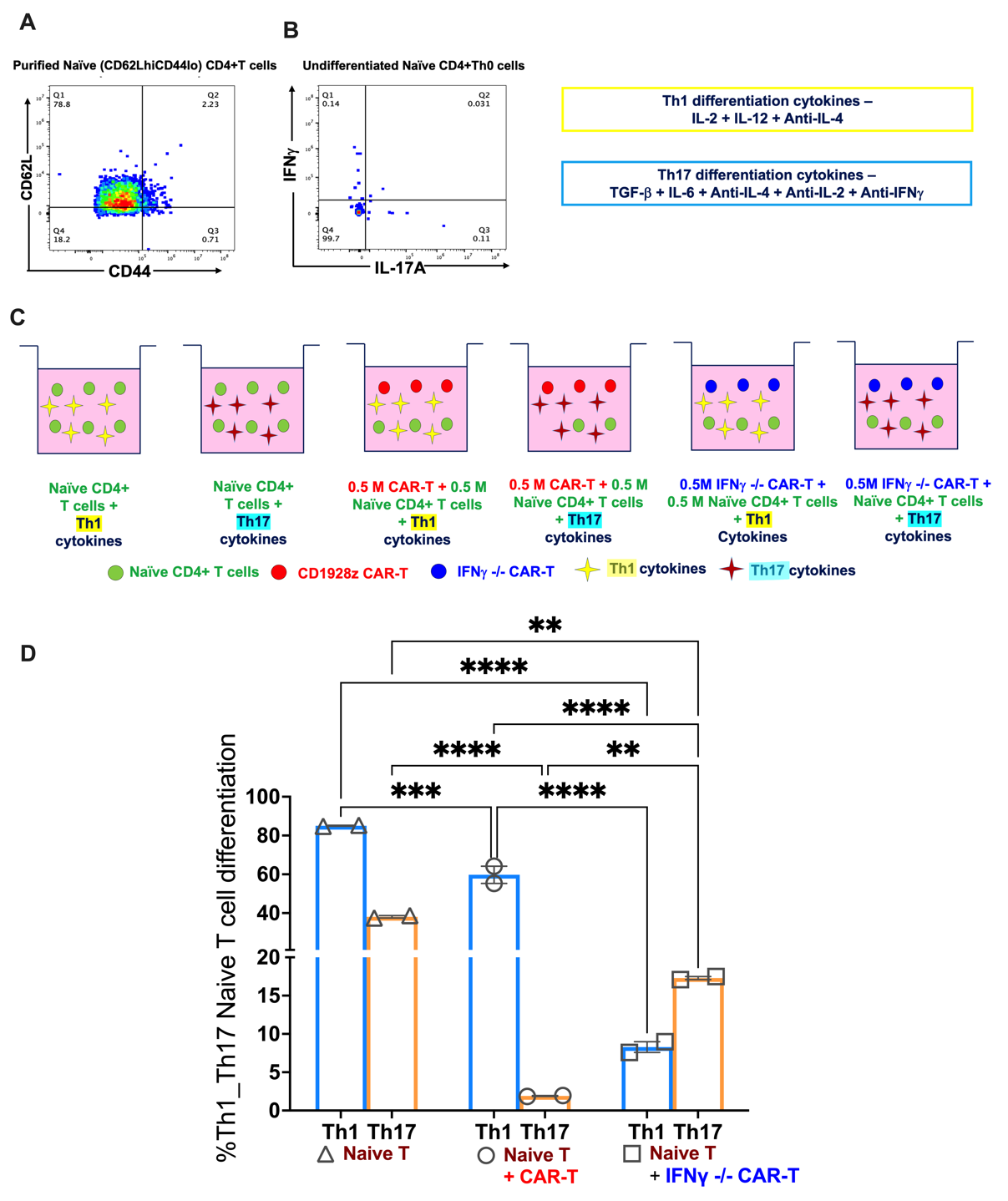
**Figure S5:** A. Naïve CD4+CD62LhiCD44lo cells were isolated (kit-based purification) and B. evaluated for Th1 cytokine IFNγ (gated as Live+ CD4+IFNγ+) and Th17 cytokine IL-17A (gated as Live+ CD4+ IL-17A+) expression. C. Purified naïve CD4+ T cells were cultured in 24 well plates and differentiated for 4 days into Th1 cells (10ng/mL IL-12, 200IU/mL IL-2 and 1 μg/mL anti-IL4) or Th17 cells (40ng/mL IL-6 , 3ng/mL TGFβ, 1 μg/mL anti-IL4, 1 μg/mL anti-IFNγ and 1 μg/mL anti-IL-2). The differentiating cells were co-cultured alone or in combination with 0.5M 1928ζ GFP CAR-T or IFNγ-/- CAR-T cells. Data is represented from a single experiment (N=2 per condition). D. Bar plots represent the percentage of naïve T cells that differentiated into Th1 (Blue bar plot gated as Live+ CD4+IFNγ+) or Th17 (Yellow bar plot gated as Live+ CD4+ IL-17A+) cells in presence or absence of CAR-T or IFNγ-/- CAR- T cells. Error bars represent SEM. P values *P < .05, **P < .01, and ***P < .001 were considered significant. p values for bar plot D was generated using Tukey’s multiple comparison test (using Two way ANOVA).

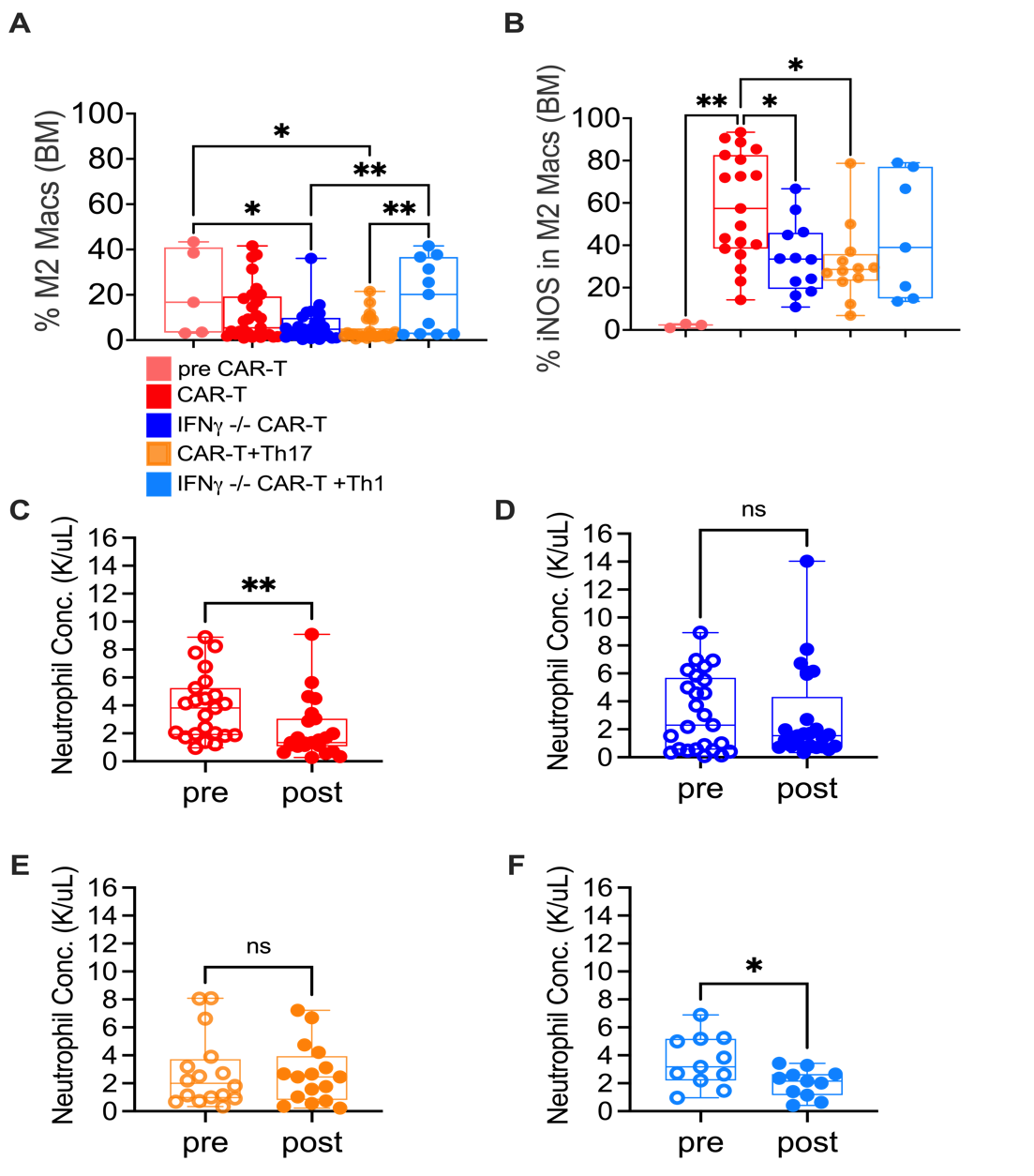
**Figure S6:** A. Arginase1+ M2-like Macrophages (gated as Live+ CD45+CD11b+F4/80+CD11c-CD206+Arginase1+ BMMCs), represented as a percent (%) of CD45+ cells were compared in mice alive at week 4 from Group 1 (N=29), Group 2 (N=28), Group 4 (N=26), and Group 5 (N=10) from Figure 5. Data are pooled from 3 independent experiments. B. iNOS+ cells in M2-like macrophages (gated as Live+ CD45+CD11b+F4/80+CD11c-CD206+Arginase1+ iNOS+ BMMCs), represented as a percent (%) of Arginase+ cells were compared in a subset of mice from Groups 1(N=19), 2 (N=12), 4 (N=12), and 5 (N=7) from Figure 5. Data are pooled from 2 independent experiments. C– F Comparison of circulating neutrophil levels from tumor bearing KO mice in Figure 5 treated with CAR-T cells with or without adoptively transferred Th17 (C, E) or IFNγ -/- CAR-T cells with or without Th1 (D, F) by CBC profiling. Pre and post neutrophil levels are paired values taken from a subset of mice alive at week 4 (Group 1 N=23, Group 2 N=25, Group 4 N= 16 and Group 5 N=11). Error bars represent SEM. P values *P < .05, **P < .01, and ***P < .001 were considered significant. P values for bar plots for M2-like macrophages in A were generated using Tukey’s multiple comparison test (using One way ANOVA). P values for neutrophil concentration plots B-E were generated using paired t test. Data for plots A-E are pooled from 3 independently performed experiments.

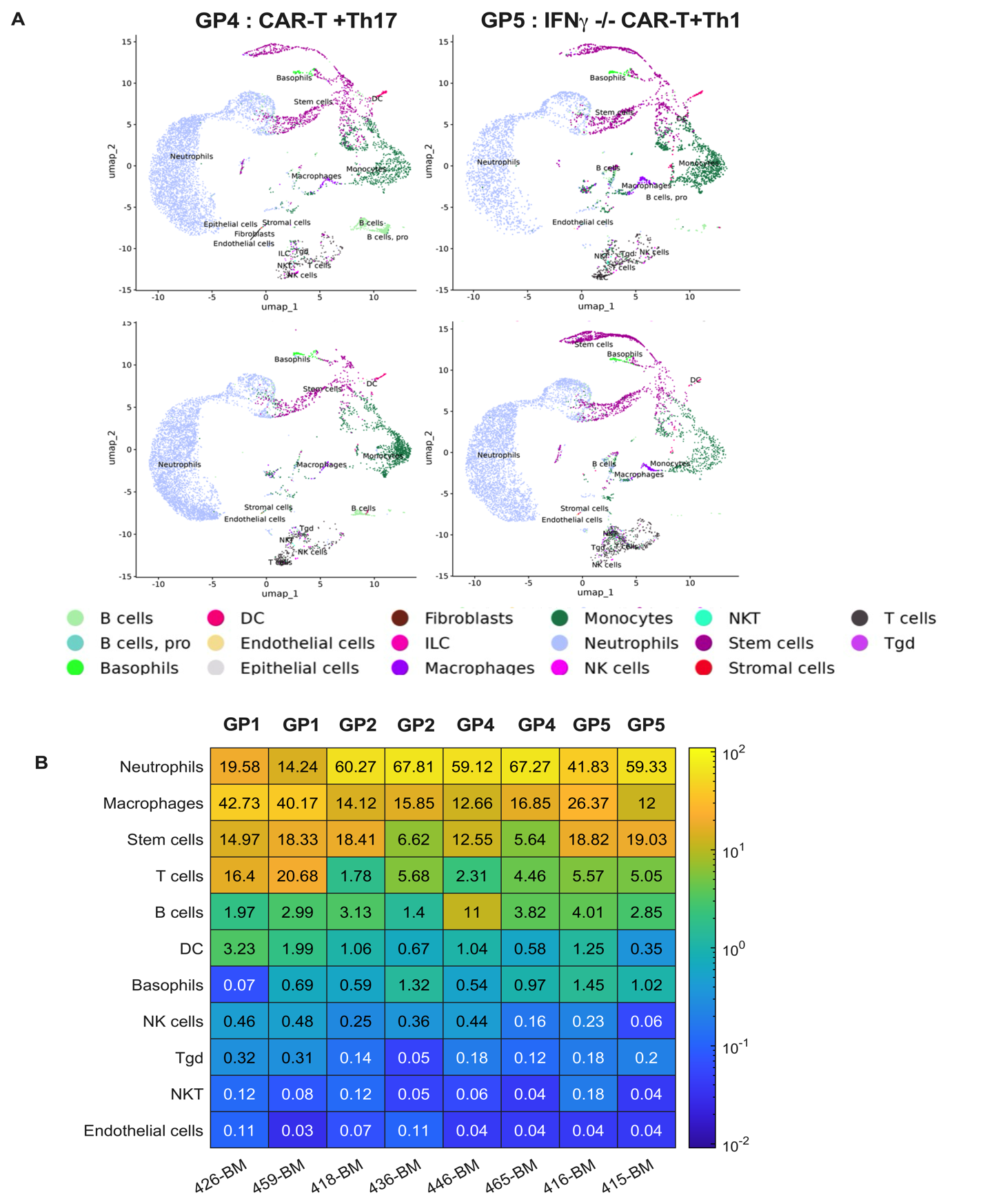
**Figure S7**: A. UMAPs of individual mice per Group (GP) 4 (CAR-T + Th17) and GP 5 (IFNg-/- CAR-T + Th1) showcasing all cells present in their bone marrow. B. Heat maps represent the frequency of various immune cells as well as stem and endothelial cells per sample in GP 1, 2, 4, and 5 described in Fig 6.

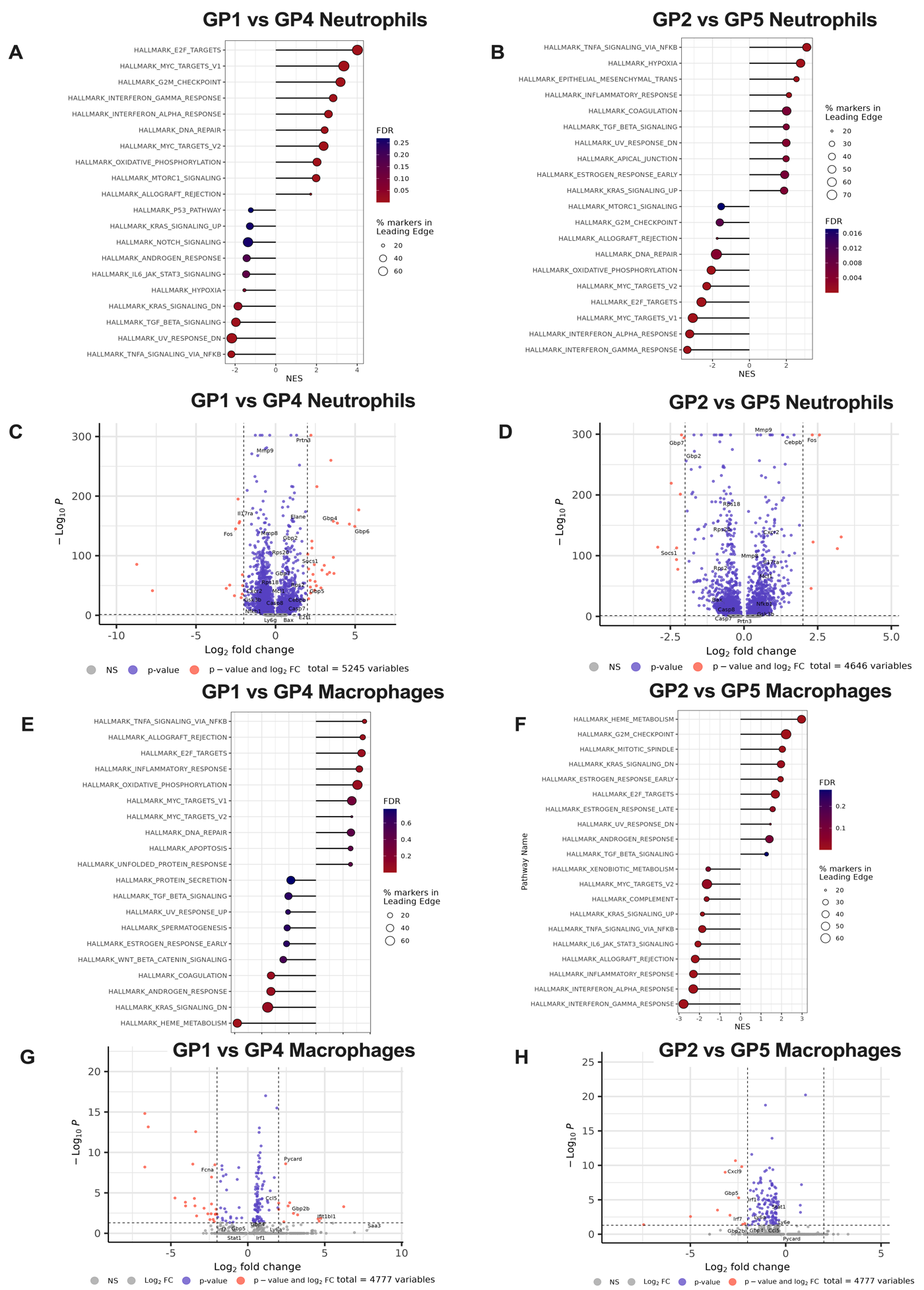

Figure S8: A-D. Comparison of gene set enrichment analysis showing hallmark pathways and respective volcano plots showing differentially expressed genes in Neutrophils from GP 1 vs GP 4 (A,C) and GP 2 vs GP 5 (B,D). E-H. Comparison of gene set enrichment analysis showing hallmark pathways and respective volcano plots showing differentially expressed genes in Macrophages from GP 1 vs GP 4 (E,G) and GP 2 vs GP 5 (F,H).

| Patient # | Disease & Diagnosis | Age (years) | Gender | CAR-T product | Best response |
| --- | --- | --- | --- | --- | --- |
| 1 | NHL, Mantle | 48 Years | Male | Brexucabtagene autoleucel | CR |
| 2 | NHL, DLBC ABC | 68 Years | Male | Axicabtagene ciloleucel | CR |
| 3 | NHL, DLBC GCB | 42 Years | Male | Lisocabtagene maraleucel | NE |
| 4 | NHL, DLBC ABC | 58 Years | Male | Lisocabtagene maraleucel | CR |
| 5 | MM, None, Lambda, IIIA | 40 Years | Male | Idecabtagene vicleucel | CR |
| 6 | MM, kappa FLC | 62 Years | Male | Idecabtagene vicleucel | VGPR |
| 7 | NHL, Mantle | 61 Years | Male | Brexucabtagene autoleucel | CR |
| 8 | NHL, DLBC ABC | 72 Years | Male | Lisocabtagene maraleucel | CR |
| 9 | NHL, DLBC GCB | 59 Years | Male | Axicabtagene ciloleucel | CR |
| 10 | NHL, Mantle | 70 Years | Male | Brexucabtagene autoleucel | CR |
| 11 | NHL, Mantle | 50 Years | Male | Brexucabtagene autoleucel | PROG |
| 12 | MM, IgG, Kappa, IIIA | 66 Years | Female | Idecabtagene vicleucel | PR |
| 13 | MM, IgD, Lambda, IIIA | 49 Years | Female | Idecabtagene vicleucel | PROG |
| 14 | MM, IgG, Kappa, IIIA | 68 Years | Male | Ciltacabtagene autoleucel | VGPR |
| 15 | MM, IgG, Lambda, IIA | 42 Years | Female | Idecabtagene vicleucel | PROG |
| 16 | MM, IgG, Kappa, IIIA | 74 Years | Male | Idecabtagene vicleucel | PROG |
| 17 | NHL, DLBC GCB | 60 Years | Male | Tisagenlecleucel | PROG |
| 18 | NHL, DLBC GCB | 36 Years | Female | Tisagenlecleucel | PROG |
| 19 | NHL, Mantle | 56 Years | Male | Brexucabtagene autoleucel | CR |
| 20 | MM, IgG, Kappa, IIIA | 60 Years | Female | Idecabtagene vicleucel | PROG |
| 21 | NHL, BcellNOS | 73 Years | Male | Lisocabtagene maraleucel | PR |
| 22 | MM, Non-sec, IIIA | 67 Years | Female | Idecabtagene vicleucel | PROG |
| 23 | NHL, DLBC ABC | 69 Years | Male | Axicabtagene ciloleucel | CR |
| 24 | NHL, DLBC NOS | 49 Years | Male | Tisagenlecleucel | PROG |
| 25 | NHL, DLBC ABC | 52 Years | Female | Tisagenlecleucel | PROG |
| 26 | NHL, DLBC ABC | 74 Years | Male | Tisagenlecleucel | CR |
| 27 | NHL, DLBC GCB | 48 Years | Male | Tisagenlecleucel | PROG |

Supplementary Table S1: Baseline demographic and clinical characteristics per patient.

NHL:non-Hodgkin lymphoma; MM:Multiple myeloma; DLBCL GCB: Germinal Center B-Cell-Like Diffuse Large B-Cell Lymphoma; DLBC ABC: Activated B-cell-like diffuse large B-cell lymphoma.

CR:complete response; PR:partial response; PROG:progression; VGPR:very good partial response.

| Patient # | Severity-based neutropenia [69] | Recovery-based neutropenia  [68] | CRS | Max Grade CRS | Risk based group classification |
| --- | --- | --- | --- | --- | --- |
| 1 | Prolonged neutropenia | Intermittent recovery | yes | 1 | Low-grade-CRS-Neutropenia |
| 2 | Prolonged neutropenia | Intermittent recovery | no | 0 | No cooccurence |
| 3 | None | Quick recovery | yes | 1 | No cooccurence |
| 4 | Profound neutropenia | Quick recovery | no | 0 | No cooccurence |
| 5 | Prolonged neutropenia | Intermittent recovery | yes | 1 | Low-grade-CRS-Neutropenia |
| 6 | Protracted + Prolonged | Intermittent recovery | yes | 1 | Low-grade-CRS-Neutropenia |
| 7 | None | Quick recovery | yes | 2 | No cooccurence |
| 8 | None | Quick recovery | no | 0 | No cooccurence |
| 9 | Prolonged neutropenia | Intermittent recovery | no | 0 | No cooccurence |
| 10 | Prolonged neutropenia | Intermittent recovery | yes | 2 | High-grade-CRS-Neutropenia |
| 11 | None | Quick recovery | no | 0 | No cooccurence |
| 12 | Profound + Protracted + Prolonged | Aplastic neutropenia | yes | 2 | High-grade-CRS-Neutropenia |
| 13 | None | Quick recovery | no | 0 | No cooccurence |
| 14 | Prolonged + profound | Intermittent recovery | yes | 3 | High-grade-CRS-Neutropenia |
| 15 | Prolonged neutropenia | Not available | no | 0 | No cooccurence |
| 16 | Protracted + Prolonged | Intermittent recovery | no | 0 | No cooccurence |
| 17 | None | Quick recovery | no | 0 | No cooccurence |
| 18 | Prolonged neutropenia | Intermittent recovery | yes | 2 | High-grade-CRS-Neutropenia |
| 19 | Protracted neutropenia | Intermittent recovery | yes | 2 | Low-grade-CRS-Neutropenia |
| 20 | Protracted + Profound | Quick recovery | yes | 2 | No cooccurence |
| 21 | Prolonged neutropenia | Intermittent recovery | no | 0 | No cooccurence |
| 22 | Profound + Protracted + Prolonged | Aplastic neutropenia | yes | 1 | Low-grade-CRS-Neutropenia |
| 23 | Prolonged neutropenia | Intermittent recovery | yes | 3 | High-grade-CRS-Neutropenia |
| 24 | None | Quick recovery | no | 0 | No cooccurence |
| 25 | None | Quick recovery | no | 0 | No cooccurence |
| 26 | Protracted neutropenia | Quick recovery | no | 0 | No cooccurence |
| 27 | Prolonged neutropenia | Intermittent recovery | yes | 1 | Low-grade-CRS-Neutropenia |

Supplementary Table S2: Patient classification based on risk of developing CRS and neutropenia
